# Readaptation of mesenchymal stem cells to high stiffness and oxygen environments modulate the extracellular matrix

**DOI:** 10.1101/2024.08.26.609692

**Authors:** Inês Caramelo, Catarina Domingues, Vera M. Mendes, Sandra I. Anjo, Margarida Geraldo, Carla M. P. Cardoso, Mário Grãos, Bruno Manadas

**Affiliations:** CNC-Center for Neuroscience and Cell Biology, University of Coimbra, 3004-504 Coimbra, Portugal; PhD Programme in Experimental Biology and Biomedicine, Institute for Interdisciplinary Research (IIIUC), University of Coimbra, Casa Costa Alemão, 3030-789 Coimbra, Portugal; CIBB - Centre for Innovative Biomedicine and Biotechnology, University of Coimbra; Institute for Interdisciplinary Research, University of Coimbra (IIIUC), 3030-789 Coimbra, Portugal; Stemlab, S.A. (Crioestaminal), 3060-197 Cantanhede, Portugal; Biocant, Technology Transfer Association, 3060-197 Cantanhede, Portugal

**Keywords:** Mesenchymal stem cells, Extracellular matrix, Secretome, Mechanomodulation, Physioxia, Proteomics

## Abstract

The therapeutic potential of mesenchymal stem cells (MSCs) has been explored over the past decades due to their ability to modulate the microenvironment through paracrine signaling. Consequently, the secretome of MCSs has emerged as a cell-free therapy rather than a cell therapy, offering the advantages of being readily commercialized as an off-the-shelf product without immunogenicity compatibility issues. As a result, strategies to manipulate and enhance the secretory profile of MSCs’ secretome are emerging. MSCs from the Wharton’s jelly niche are accommodated to the stiffness and oxygen level found at the umbilical cord (UC), which are 2 to 5kPa (Young’s modulus) and 2.4% to 3.8% O_2_, respectively. However *in vitro* culture conditions (2-3 GPa and 18.5% O_2_) are largely different from the one observed in vivo. Here, we present a proteomic characterization of the secretome of MSCs primed (48h) or readapted (7-10 days) to soft (3kPa) (mechanomodulated) or low oxygen levels (5% O_2_) (physioxia). Maintaining MSCs on soft platforms for long periods increased the secretion of proteins associated with cell redox homeostasis, such as protein disulfide isomerases and mitochondrial proteins, while physioxia enhanced the secretion of immunomodulatory proteins. The high secretion of these proteins might confer a therapeutical advantage by favoring a regenerative environment at the injury site. Interestingly, lowering the stiffness or oxygen converged on the downregulation of several extracellular matrix proteins (ECM), particularly collagen fibrils, on primed and readapted cells. These results suggest that a massive reorganization of the extracellular space occurs upon culturing MSCs on conventional culture conditions, which may affect not only matrix stiffness but also several signaling pathways initiated at the cell membrane, such as PDGF signaling pathways (e.g., PI3K-AKT), consequently biasing stem cell fate. In conclusion, mimicking physiological culture conditions *in vitro* modulates secretome composition, which may empower its therapeutical properties by enriching proteins that promote cell survival.

**Graphical Abstract:** 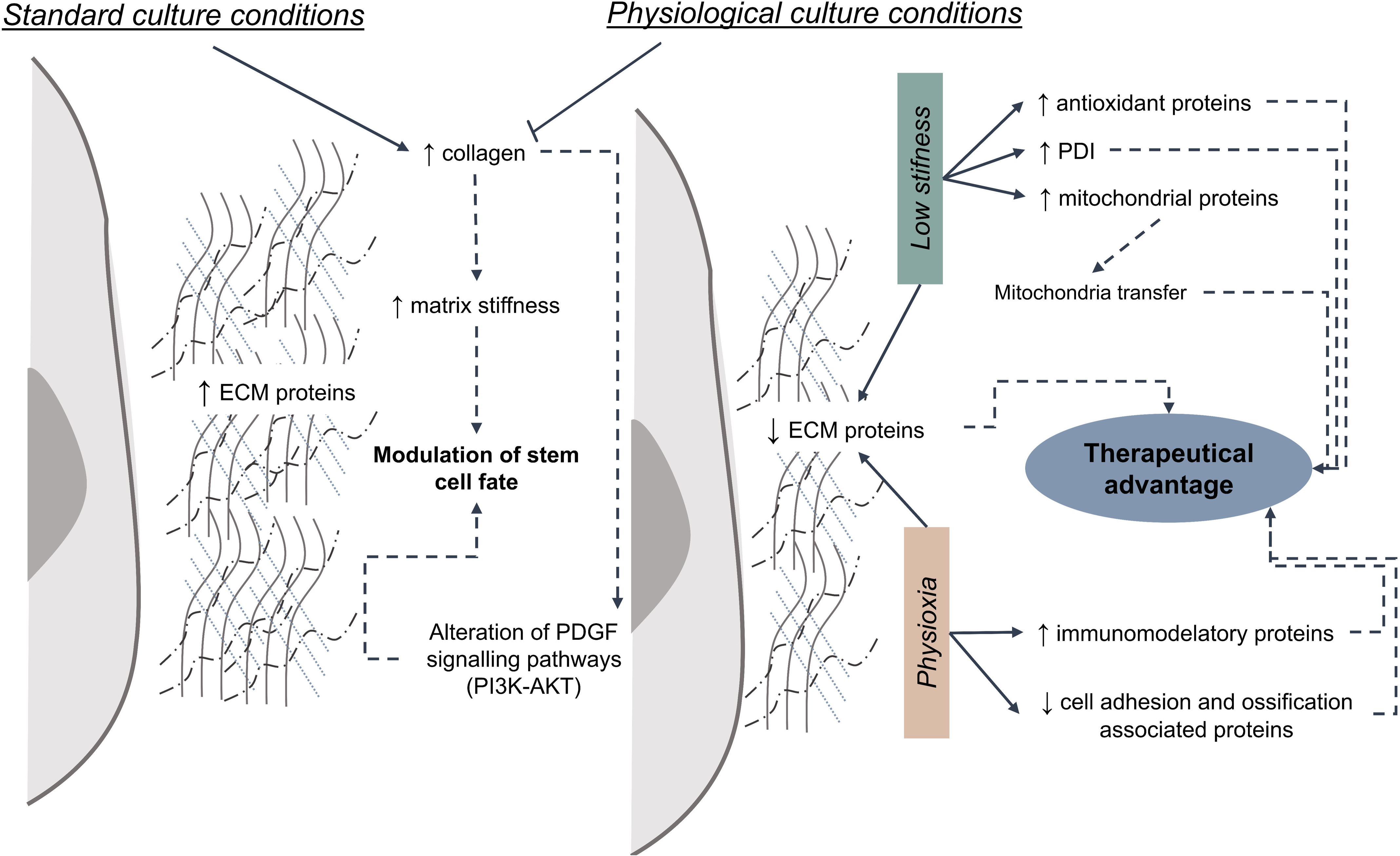

## Introduction

The therapeutic potential of mesenchymal stem cells (MSCs) has gathered significant attention in the regenerative medicine field (1). These cells exhibit the capability not only to self-renewal *in vitro* but also to differentiate into multiple lineages, making them suitable for a broad spectrum of clinical applications (2, 3). For many years, MSCs treatments comprised mainly the autologous or allogenic transplantation of cells (“cell-based therapy”) (4). While the former is described to have lower associated risks, recent clinical trials have shown that allogenic MSCs are also a safe and promising treatment approach (5, 6). Once administered, MSCs are chemoattracted to the injury site, where they promote tissue repair (1, 7). Beyond their regenerative role through the differentiation and replacement of damaged cells, MSCs are renowned for their ability to modulate the microenvironment through paracrine signaling, highlighting the increasing importance of this fluid in regenerative medicine (8). Recent studies have used the secretome of MCSs as a “cell-free therapy”. The secretome is the collection of bioactive substances secreted by cells into the extracellular space and can be categorized into two main parts: the soluble fraction, containing growth factors and cytokines, among others, and the vesicular fraction, which includes exosomes and larger vesicles (8, 9). These components can modulate different processes in the surrounding cells, such as immunomodulation, apoptosis, angiogenesis, and antioxidation (8, 10). As a result, the secretome administration is emerging as an alternative to cell therapy. This approach offers numerous advantages, particularly in terms of safety, since it avoids immunogenicity and compatibility issues and is more comparable to conventional pharmaceutical products in terms of dosage and potency (11, 12). Additionally, it can be readily commercialized as an off-the-shelf product, with extended storage lifespans without compromising potency, and is more easily scalable, thus making it financially viable. However, the secretome composition is highly dynamic and can vary depending on the MSCs source and in response to subtle different culture conditions (13, 14), which can affect consistency between batches.

Recently, strategies to manipulate and enhance the secretory profile of MSCs secretome have been explored. These approaches involve either culturing these cells on 2D or 3D hydrogels with customized biochemical and biophysical properties, using priming strategies by adding drugs and soluble factors, such as cytokines, or replicating hypoxic environments (15, 16).

MSCs can sense and respond to biophysical properties of the surrounding environment, including stiffness and tissue oxygen levels (17, 18). For instance, MSCs at the Wharton’s jelly niche respond to the stiffness and oxygen level found at the umbilical cord (UC), which is 2 to 5kPa (Young’s modulus) and 2.4% to 3.8% O_2,_ respectively (19, 20). To replicate these physiological conditions *in vitro,* we previously optimized culture conditions for MSCs on softer polydimethylsiloxane (PDMS) platforms (∼3kPa) or under controlled oxygen levels (∼5% O_2_) (21). The proteomic characterization unveiled alterations at the cellular level and a differential expression of extracellular matrix (ECM) proteins, suggesting that the secretory profile of these MSCs might also be altered in response to these physiological cues. In this study, we provide a proteomic analysis of the secretome of MSCs isolated from the umbilical cord (UC-MSCs) cultured on 3kPa platforms (mechanomodulated) and exposed to controlled levels of oxygen (5% O_2_, physioxia) for 7-10 days (readapted) or 48 hours (primed).

## Methods

### Cell culture

#### Isolation and expansion of UC-MSCs

Experiments were performed as previously described (21, 22). All the procedures were approved by the Ethics Committee of the Faculty of Medicine, University of Coimbra, Portugal (ref. CE-075/2019). Briefly, MSCs were isolated from fragments from the UC matrix (Wharton’s jelly) and expanded with proliferation medium (Minimum essential Medium-α (MEM-α) (Gibco™) supplemented with 10% (v/v) MSC-qualified fetal bovine serum (FBS) (Hyclone, GE Healthcare) (only for readapted mechanomodulated MSCs) or 5% (v/v) fibrinogen depleted human platelet lysate (HPL) (UltraGRO™, Helios) and antibiotics: 100 U/ml of Penicillin, 100 μg/ml Streptomycin and 2.5 μg/ml Amphotericin B or Antibiotic-Antimycotic (all from Gibco™). When cells were confluent, MSCs were trypsinized by adding 0.05% Trypsin-EDTA solution (Gibco™) for five minutes at 37°C, homogenized, centrifuged, and seeded at the desired density.

For standard culture conditions, UC-MSCs were kept on a humidified incubator at 37°C, with 5% CO_2_, on a tissue culture polystyrene (TCP) dish, from P2 to P4. For priming experiments, cells were seeded at 10.000/cm^2^ (P4).

#### MSCs expansion under physiological culture conditions

Two different physiological environments were created separately: low stiffness or oxygen levels. Mechanomodulated MSCs were kept under low stiffness conditions (3kPa). To create soft culture conditions, platforms of polydimethylsiloxane (PDMS) were prepared and functionalized as previously described (22). Physioxia MSCs were maintained in a low oxygen environment. For that, MSCs were cultured in an InvivO₂^®^ 400 Physoxia Workstation (Baker), with 5% O_2_, 5% CO_2_ and humidified environment, in accordance with the protocol previously described (21).

For both experimental conditions, UC-MSCs were expanded under conditions that mimic a physiological environment (3kPa platforms or 5%O_2_) for 7-10 days, as the population doubling depends on the donors, (hereafter termed “re-adapted”) or for 48h (termed “priming”). To prepare the conditioned medium (CM), UC-MSCs were washed twice with PBS, and a fresh culture medium without supplements (only antibiotics) was added. This procedure aims to remove supplemented proteins, which, owing to their high abundance, might compromise the detection of secreted proteins due to their low abundance. UC-MSCs were left to secrete for 48h (readapted) or 24h (priming).

### Proteomic analysis

#### Secretome preparation

The secretome was prepared as previously described (23). Briefly, CM was collected and centrifuged at 290×g for 5 min at 4°C. The 4μg of protein standard (MBP-GFP) was added to the same volume of secretome (24). Then, the supernatant was transferred to a low molecular weight cut-off (5kDa Vivaspin^®^ 20, Sartorius) and centrifuged at 3,000×g, at 4°C, until the desired concentration was achieved. Then, proteins were precipitated through the sequential addition of trichloroacetic acid and acetone and resuspended on sample buffer 2× (116.7 mM Tris-HCl pH 6.8, 10% (v/v) Glycerol, 3.3% (w/v) SDS, 0.31% (w/v) dithiothreitol (DTT), bromophenol blue).

For the proteomic analysis, the total volume of CM was analyzed, except for physioxia readapted secretome, where an equal amount of protein was loaded in the gel (50μg). Proteins were first alkylated by the addition of a 40% (w/v) acrylamide solution and resolved through a short-GeLC approach and stained with Coomassie Brilliant Blue G-250 (25, 26). Pools of each condition were prepared for protein identification. Concerning protein quantification, each sample was loaded individually. Gel lanes were cut and destained, and proteins were digested overnight in gel using trypsin. Finally, the peptides were extracted through the sequential addition of solutions with increasing acetonitrile concentration (30%, 50% and 98%) and 1% formic acid, dried by vacuum centrifugation at 60°C, and stored at -20°C.

#### Data acquisition

Samples were analyzed as previously described (21). Briefly, samples were resuspended in a solution of 2% acetonitrile and 0.1% formic acid and analyzed using a NanoLC™ 425 System (Eksigent) coupled to a Triple TOF™ 6600 mass spectrometer (Sciex) equipped with an electrospray ionization source (DuoSpray™ Ion Source from Sciex). Peptides were separated on a Triart C18 Capillary Column 1/32” (12 nm, 3 μm, 150 mm × 0.3 mm, YMC) and using a Triart C18 Capillary Guard Column (0.5 mm × 5 mm, 3 μm, 12 nm, YMC) at 50°C. The flow rate was set to 5 µL/min and mobile phases A and B were 5% DMSO plus 0.1% formic acid in water and 5% DMSO plus 0.1% formic acid in acetonitrile, respectively. The LC program was performed as follows: 5 - 30% of B (0 - 50 min), 30 - 98% of B (50 - 52 min), 98% of B (52-54 min), 98 - 5% of B (54 - 56 min), and 5% of B (56 - 65 min). The ionization source was operated in the positive mode set to an ion spray voltage of 5500 V, 25 psi for nebulizer gas 1 (GS1), 10 psi for nebulizer gas 2 (GS2), 25 psi for the curtain gas (CUR), and source temperature (TEM) at 100°C. For Data Dependent Acquisition (DDA) experiments, the mass spectrometer was set to scan full spectra (m/z 350-2250) for 250 ms, followed by up to 100 MS/MS scans (m/z 100-1500) with a 30 ms accumulation time, resulting in a cycle time of 3.3 s. Candidate ions with a charge state between + 1 and + 5 and counts above a minimum threshold of 10 counts per second were isolated for fragmentation, and one MS/MS spectrum was collected before adding those ions to the exclusion list for 15 s (mass spectrometer operated by Analyst^®^ TF 1.8.1, Sciex). For Data Independent Acquisition (DIA) (or Sequential Window Acquisition of all Theoretical Mass Spectra (SWATH)) experiments, the mass spectrometer was operated in a looped product ion mode and specifically tuned to a set of 42 overlapping windows, covering the precursor mass range of 350–1400 m/z. A 50 ms survey scan (m/z 350–2250) was acquired at the beginning of each cycle, and SWATH-MS/MS spectra were collected from 100 to 2250 m/z for 50 ms, resulting in a cycle time of 2.2 s.

#### Data analysis

The five fractions of each pool were acquired individually in DDA mode. To generate the ion library, all files were combined and searched against the reviewed Human (Swiss-Prot) database (downloaded on 21^st^ October 2021 (for re-adapted MSCs experiments) and 16^th^ May 2022 (for priming) using the ProteinPilot™ software (v5.0, Sciex). An independent False Discovery Rate (FDR) analysis was performed using the target-decoy approach provided by the software to assess the quality of the identifications. Concerning protein relative quantification, samples were acquired on DIA mode (SWATH) and analyzed individually. Data processing was performed using the SWATH™ plug-in for PeakView™ (v2.0.01, Sciex), configured to select up to 15 peptides per protein and five transitions per peptide. To avoid the interference of FBS contaminants on long exposure mechanomodulation experimental setup, a sequence list of highly abundant fetal bovine serum proteins was added to the database (27). These proteins were removed from the analysis.

Peptides were only quantified if they had an FDR below 1% in at least one-third of the samples per condition and were normalized to the total intensity of the sample. The MS data have been deposited to the ProteomeXchange Consortium via the PRIDE partner repository with the dataset identifier PXD052907. *Credentials for reviewers only: Username: Password: sXN2pzxkN24C* MetaboAnalyst 5.0 online platform and GraphPad Prism 9 were used to perform the statistical analyses (28). Both univariate (t-test or Mann-Whitney or Kruskal-Wallis tests) and multivariate (Partial least squares-discriminant analysis (PLS-DA)) analyses were used to identify proteins of interest. Hits were selected if they had a VIP>1, p<0.05, or both, and fold change (FC) above 1.5 or below 0.67 (|Log_1.5_FC|>1) for further analysis. FunRich was used to perform a Gene Ontology (GO) enrichment analysis (29), while pathways were mapped using the Reactome database (30). Pathways were searched on KEGG database (31). Violin plots were created on GraphPad Prism 9 and Venn diagrams were generated on DeepVenn (32).

## Results

### Readapting UC-MSCs to low stiffness or oxygen environments modulates their secretory profile

MSCs can act through paracrine signaling to promote tissue regeneration at the site of injury; therefore, unveiling the composition of their secretome has emerged as a hot topic in this research field. Previous data indicated that culturing MSCs on lower stiffness platforms modulated the cellular proteome, including components of the extracellular matrix (21, 22). Therefore, it was hypothesized that the secretome composition might also be altered in response to mechanical cues. Hence, UC-MSCs were expanded on 3kPa platforms (P2-P4), and the CM was collected and subjected to a proteomic analysis. From the 1007 proteins identified, 705 proteins were quantified with confidence and subjected to statistical analysis to identify significantly secreted proteins under a low stiffness environment, compared to standard culture conditions. The PLS-DA scores plot showed no overlap between the two experimental groups (Figure 1A). Additionally, using a univariate approach, several proteins were identified as more or less abundant (Figure 1B). Next, the 193 significant proteins (VIP>1, p<0.05, or both, and |Log_1.5_FC|>1) were subjected to a GO analysis. Looking at the biological processes, upregulated proteins are mainly associated with carbohydrate metabolism, cell redox homeostasis, and protein folding processes (Figure 1C). In contrast, proteins associated with translation and extracellular matrix organization cell adhesion decrease in the CM of UC-MSCs kept in soft platforms.

**Figure 1.**
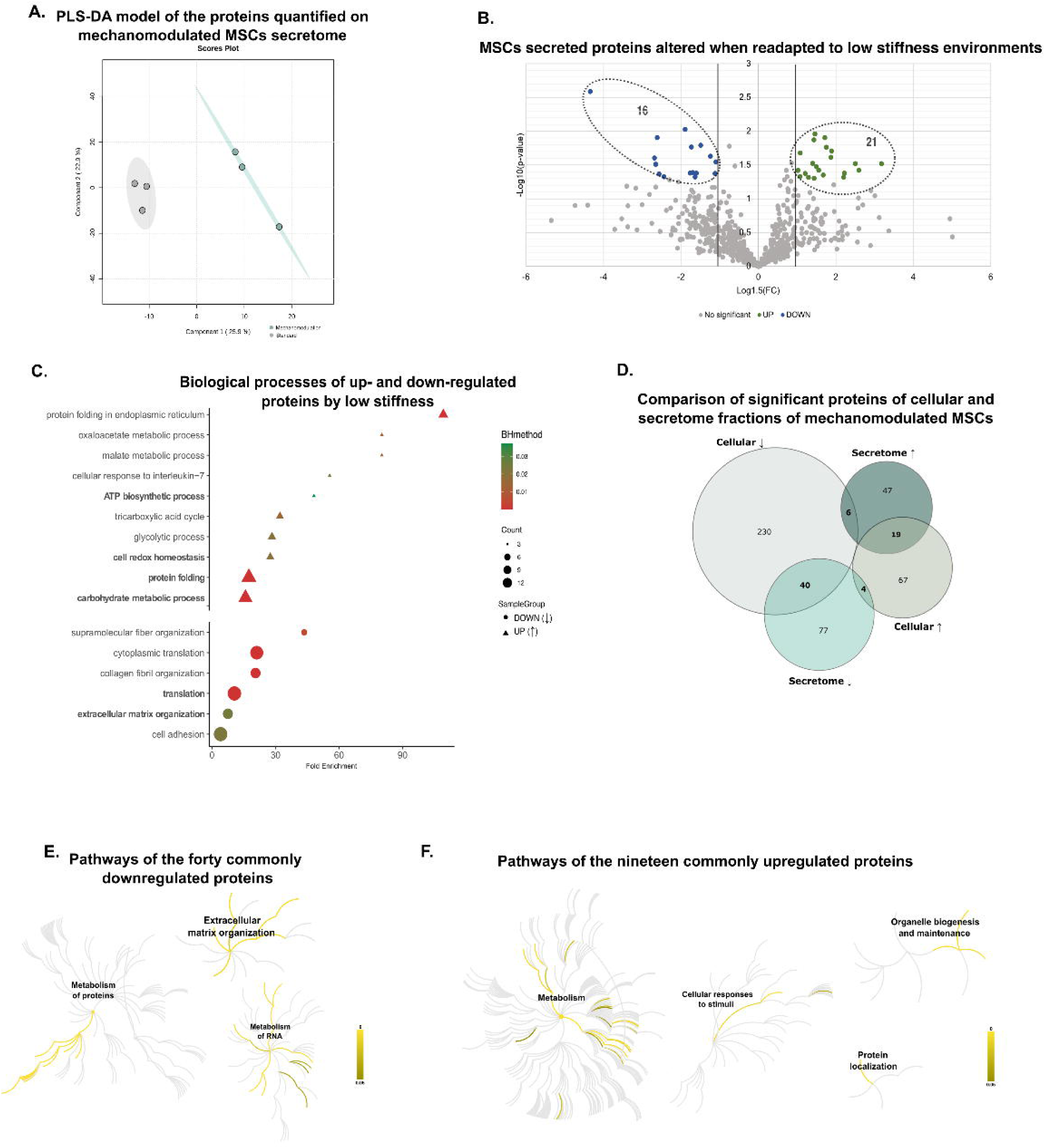
Proteins involved in carbohydrate, RNA, and protein metabolism and ECM are differentially secreted by MSCs readapted to low stiffness conditions. All quantified proteins on both standard (grey) and mechanomodulation (green) groups were used to generate a PLS-DA scores plot (A). The statistically different proteins (t-test, p<0.05; |Log_1.5_FC|>1) are highlighted on the Volcano Plot (upregulated in green; downregulated in blue) (B). In total, 121 and 72 significant proteins were found to be down-(↓) or up-(i) regulated, respectively (VIP>1, p<0.05, or both; |Log_1.5_FC|>1). These proteins were subjected to a GO analysis. Significant biological processes (GO) are represented (circles for downregulated and triangles for upregulated proteins) (C). Next, differentially expressed proteins of the proteome and secretome were compared (D). Common proteins shared by both fractions were then analyzed. The most significant Reactome pathways of the commonly forty down- and nineteen up-regulated proteins are highlighted in yellow (E and F, respectively; Supplementary Tables 1 and 2). Abbreviations: Benjamini-Hochberg method (BH).

To evaluate if the secretory profile rate was being altered, UC-MSCs, the proteome of both the cellular (previously published (21)) and secretome fractions was compared, where 11.2% and 13.3% of the proteins where found to be up- and down-regulated on both fractions, respectively (Figure 1D). Proteins found to be decreased both on the cellular proteome and secretome were mainly involved in the protein metabolism (e.g., translation), RNA processes (e.g., rRNA processing), and ECM organization (e.g., collagen formation) (Figure 1E, Supplementary Table 1). On the other hand, cellular carbohydrate metabolism also appears to be the main mechanism altered when considering upregulated proteins since the levels of several proteins associated with the citric acid (TCA) cycle, respiratory electron transport, and mitochondrial biogenesis increased when cells were readapted to low stiffness environments (Figure 1F, Supplementary Table 2). Besides, proteins commonly upregulated were found to be associated with cell detoxification of reactive oxygen species (ROS) processes Only a few proteins (10 out of 490 proteins) exhibit an opposite profile when comparing the cellular proteome and secretome, suggesting that these proteins might either be more secreted to the extracellular environment or accumulate inside the cell, decreasing the secretion in response to the physiological environment (Supplementary Figure 1A-B, respectively).

The overall results suggest that when UC-MSCs are exposed to prolonged periods in culture conditions with low stiffness, the composition of the secretome changes. This may be due to an alteration in protein abundance inside the cell and the maintenance of secretion rates or a result of regulation of the secretion rate in response to the extracellular environment.

Previous data reported that UC-MSCs cellular proteome was altered when cells were maintained under physiological oxygen levels, particularly interfering with protein metabolism processes (21). To investigate if changes in these processes impacted UC-MSCs’ secretory profile, the CM of UC-MSCs readapted to physioxia (5% oxygen) was explored. After expansion under low oxygen levels (7 to 10 days), the CM of UC-MSCs was collected for proteome analysis using the above-mentioned pipeline. The generated library comprised 1535 proteins identified with confidence, from which 450 proteins were quantified with confidence (SWATH-MS). To identify interesting targets, all quantified proteins underwent a univariate and multivariate analysis approach. The two experimental groups do not intersect on the PLS-DA scores plot (Figure 2A). In addition, the volcano plot highlights significant up- and downregulated proteins (Figure 2B). To disclose the biological processes in which proteins of interest play a role, these proteins were subjected to a GO analysis. Data revealed that upregulated proteins are mainly associated with fibrinolysis and innate and adaptative immune responses. In contrast, proteins of the extracellular matrix and involved in cell adhesion processes appear to be downregulated (Figure 2C, Supplementary Table 3).

**Figure 2.**
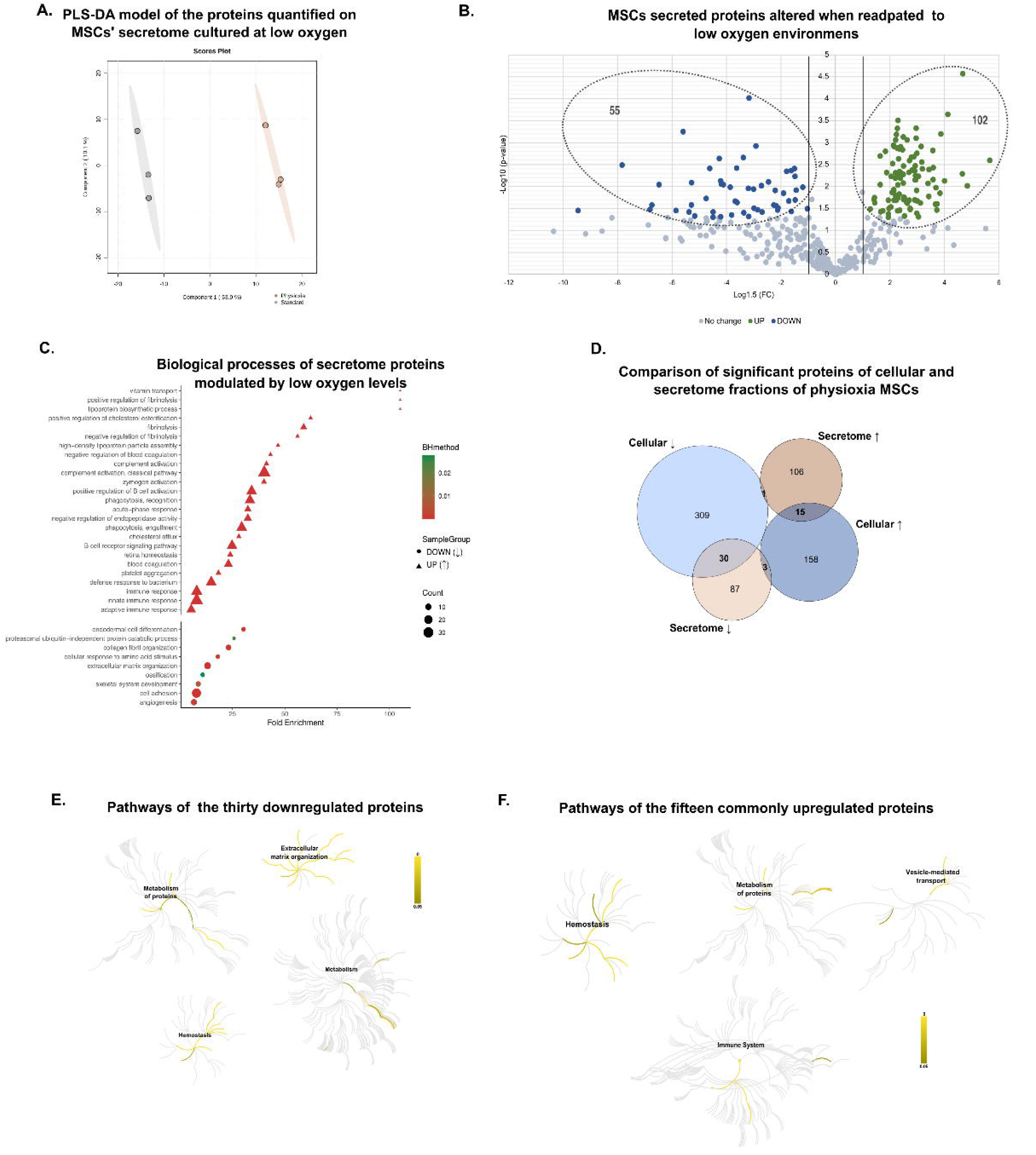
Readapting MSCs to low oxygen environments modulates the secretion of proteins involved in metabolism, the immune system, and ECM. The PLS-DA scores plot (A) represents all quantified proteins. The Volcano Plot highlights significant proteins up- (i) (green) and down- (↓) (blue) regulated (t-test, p<0.05; |Log1.5FC|>1) (B). Data analysis revealed that 120 and 122 proteins were down- (↓) and up- (i) regulated, respectively (VIP>1, p<0.05, or both; |Log_1.5_FC|>1). (C) Biological processes (GO) of these proteins (BH method, p<0.001; all significant hits are available in Supplementary Table 3). The statistically altered proteins for the secretome and previously published cellular fraction (21) were compared (D). Proteins highlighted in the intersections were further explored. Significant Reactome pathways of the thirty down- and fifteen up-regulated proteins shared by both screenings are represented in yellow (E and F, respectively; Supplementary Tables 4 and 5). Abbreviations: Benjamini-Hochberg method (BH).

Next, the protein levels of the cellular proteome and secretome were compared to identify trends in secretion profiles modulated by physioxia oxygen levels culture conditions, with 5.3% of the proteins found to be commonly increased on both fractions, while 7.0% were decreased (Figure 2D). Proteins associated with extracellular matrix organization, mainly collagen, elastic fibers, and fibronectin components, are downregulated both in the intra and extracellular environment (Figure 2E, Supplementary Table 4). Moreover, the metabolism of glycosaminoglycans and post-translational modifications of proteins, including phosphorylation and calnexin/calreticulin cycle, were also described to be downregulated. Conversely, proteins associated with innate immune responses were increased in both proteome and secretome, as well as proteins involved in forming the fibrin clot (hemostasis) and scavenging the heme from the plasma (Figure 2F, Supplementary Table 5). Finally, four proteins lyase display inverse tendencies on both fractions, suggesting that the cellular levels of these proteins might be influenced by increased or decreased secretory rates or altered vesicle transport dynamics, including internalization processes (Supplementary Figure 1C-D).

### Extracellular matrix (ECM) proteins play a key role in the adaptation of UC-MSCs to physiological environments

Data analysis revealed that mimicking the physiological stiffness and oxygen levels environment *in vitro* led to a modulation of the extracellular matrix organization, cell adhesion, and the metabolism of proteins. To explore common mechanisms shared by these physiological-like environments, statistically different proteins previously described for both screenings were compared (Figure 3A). Only ATP synthase subunit alpha (mitochondrial) was found to be upregulated in both conditions (Supplementary Figure 1E). However, the levels of Talin-1, a protein involved in cell-matrix interaction, increased on physioxia-modulated secretome and decreased on mechanomodulated secretome (Supplementary Figure 1F). The majority of common proteins was found to be downregulated when MSCs were readapted to low oxygen or stiffness environments. The role of these proteins was further explored using Reactome database. Results indicate that these proteins are associated with the extracellular matrix organization, namely collagen, and proteins involved in the cross-linking of these fibrils, as peroxidasin homolog, and proteoglycans, as well as some proteins involved in MET (Hepatocyte growth factor receptor) or PDGF (Platelet-Derived Growth Factor) signaling pathways (Figure 3B, D, Supplementary Table 6). In addition, the ten proteins with inverse tendencies in both physiological environments were found to be involved in glucose metabolism and protein post-translational modifications, particularly affecting the calnexin/calreticulin cycle (Figure 3F and Supplementary Figure 1G, Supplementary Table 7). It can be concluded that similarly to what has been observed for the cellular proteome (33), the two different physiological environments culminate in the diminished secretion of common proteins, particularly ECM proteins.

**Figure 3.**
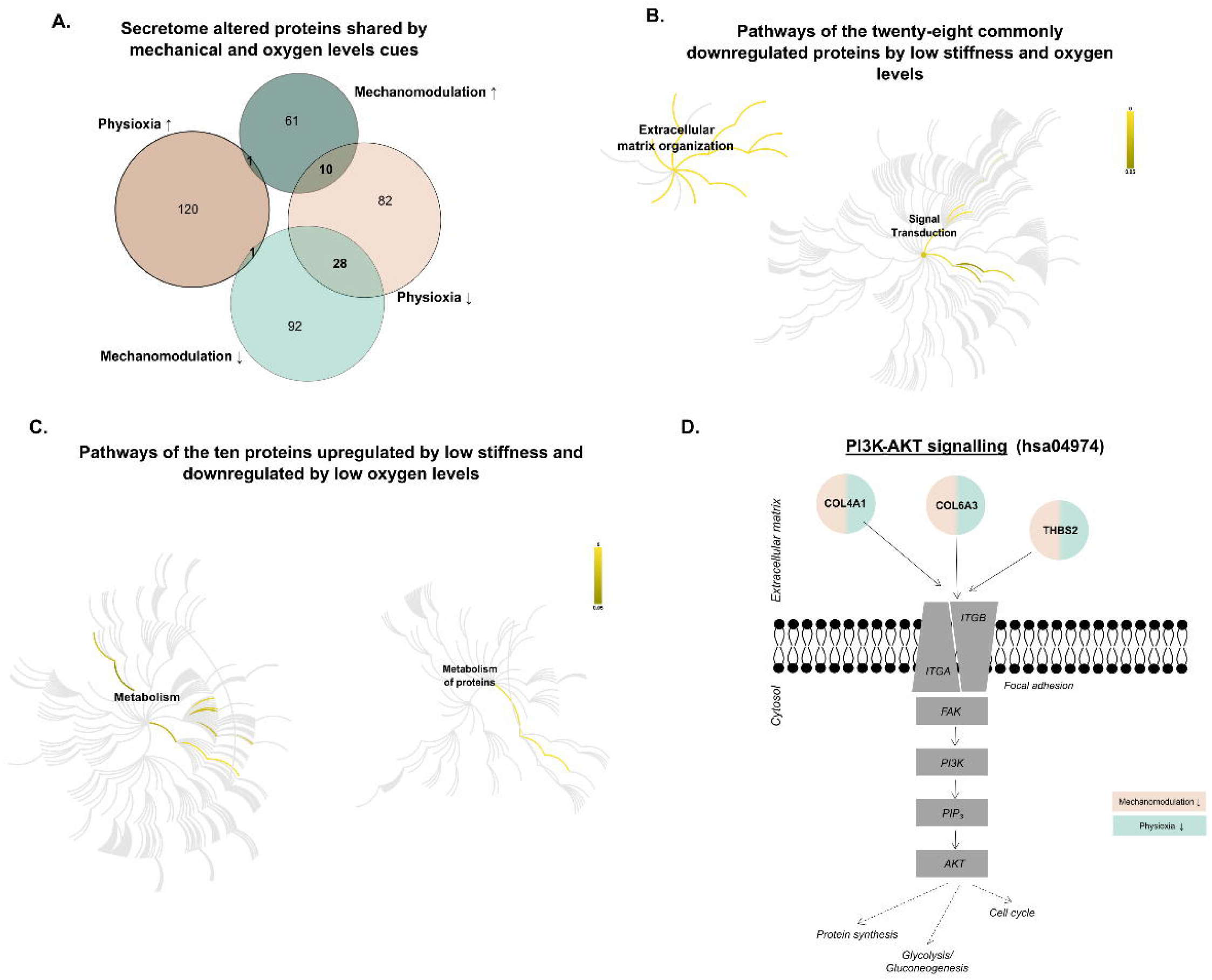
UC-MSCs kept under low stiffness or oxygen levels conditions commonly reorganize the secretion of ECM proteins. Significant proteins previously selected (VIP>1, p<0.05, or both; |Log_1.5_FC|>1) from each screening were compared and are represented on the Venn diagram (A). Proteins shared by both experimental conditions were further analyzed. Significant Reactome pathways of the twenty-eight proteins commonly downregulated under physiological conditions (B) and the ten proteins upregulated by mechanomodulation and downregulated by physioxia (C) are represented in yellow (details can be consulted in Supplementary Tables 6 and 7). In detail, the proteins involved in the PDGF signaling (that can activate PI3K/AKT signaling pathway) (D) are highlighted in circles, according to the Kegg Pathway database (grey: proteins that belong to the pathway; light green: downregulated on mechanomodulation; light brown: downregulated on physioxia). Abbreviations: COL4A1: collagen type IV alpha 1 chain; COL6A3: collagen type VI alpha 3 chain; THBS2: thrombospondin 2; ITGA: integrin subunit alpha; ITGB: integrin beta; FAK: protein tyrosine kinase 2; PI3K: phosphatidylinositol-4,5-bisphosphate 3-kinase catalytic subunit alpha; PIP_3_: phosphatidylinositol-3, 4, 5-triphosphate; AKT: AKT serine/threonine kinase 3)

Expanding UC-MSCs under standard culture conditions for long periods (7-10 days) does not impair UC-MSCs’ ability to adapt to lower stiffness and oxygen culture conditions, as previously published (21). Therefore, it was hypothesized that the secretory profile of MSCs was able to adapt to physiological environments for a short period after being expanded at standard conditions – priming. Briefly, UC-MSCs were expanded under standard culture conditions until passage 3, after which cells were primed for 48h, using low stiffness platforms (3kPa) or a physioxia environment (5% O_2_) or maintained in standard conditions. Then, primed UC-MSCs secretome was collected, and the proteome was analyzed as previously. This resulted in the identification of 557 proteins, from which 413 were quantified with confidence in all experimental groups. The cluster analysis and the PCA scores plot demonstrate a closer relationship between the standard and physioxia secretome despite the separation of the three groups on the PLS-DA model (Figure 4A and Supplementary Figure 2A-B, respectively). Next, the GO analysis revealed that keeping cells on low-stiffness platforms decreases the secretion of proteins involved in cell adhesion, more specifically, various types of collagen, as described in long-exposure experiments (Figure 4B, 1C, and Supplementary Figure 2D). In parallel, physioxia priming upregulates proteins associated with immunomodulatory responses, as observed in long-exposure experiments Figure 4B and Supplementary Figure 2D). Interestingly, the extracellular matrix components, particularly the collagen fibril organization, were found to be downregulated upon both physiological priming, consistent with the secretory profile of readapted MSCs. Together, this data suggests that the extracellular matrix is a key player in adapting UC-MSCs to standard culture conditions, possibly leading cells to increase the expression and secretion of these components.

**Figure 4.**
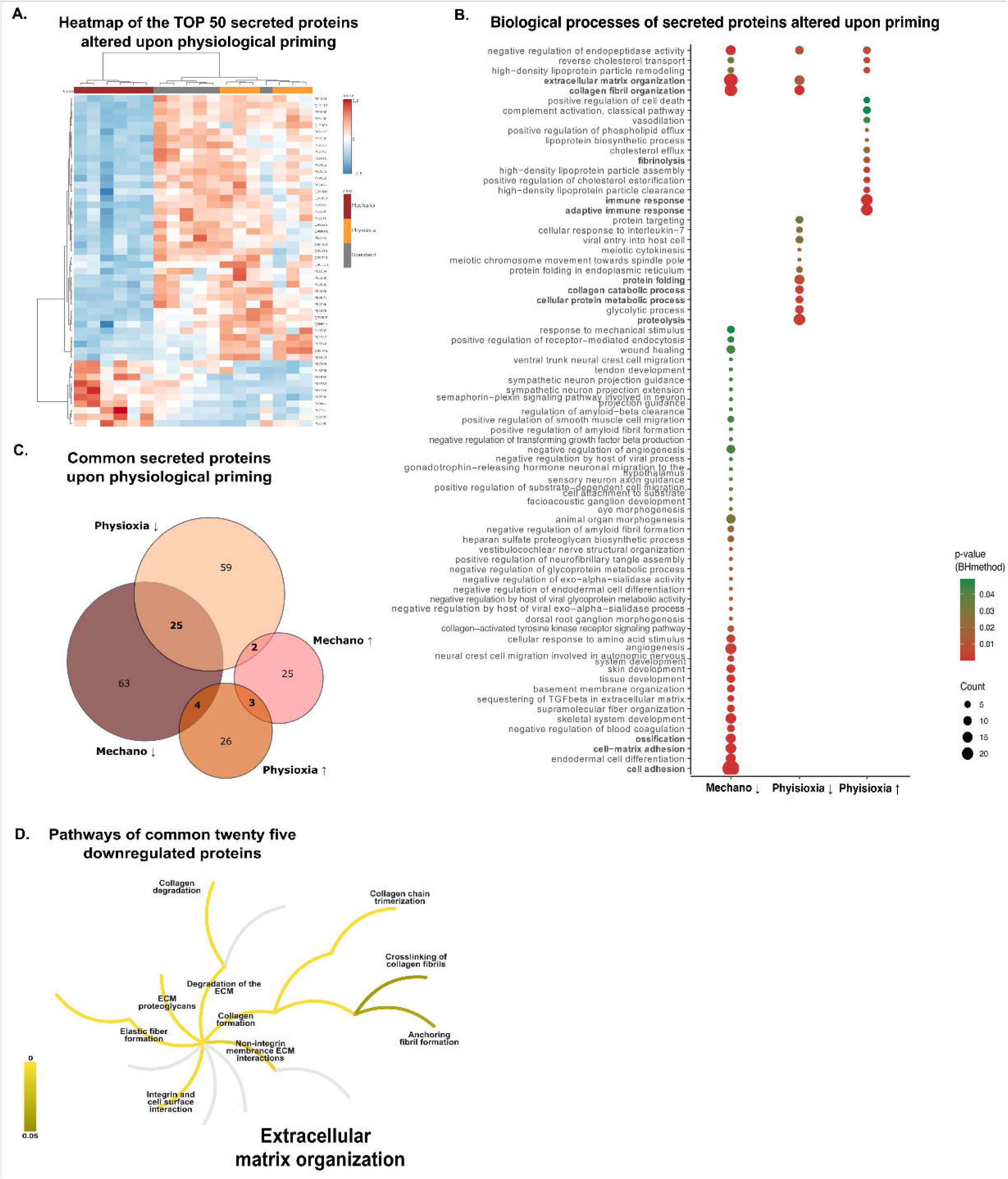
Physiological priming decreases the secretion of ECM proteins. The top 50 proteins (VIP>1, Supplementary Figure 2B) quantified with confidence on the three experimental groups: Standard (grey), Mechanomodulation (red) and Physioxia (yellow) are represented on the heatmap (with hierarchical clustering) (A) (protein names on Supplementary Table 8). The mechanomodulated secretome comprised 30 up-regulated (i) proteins and 92 down-regulated (↓) proteins. Lowering the oxygen levels increased the secreted levels of 33 proteins and decreased the levels of 86 proteins. The significant biological processes (GO) of the filtered proteins (VIP>1, p<0.05, or both; |Log1.5FC|>1) from each experimental condition are highlighted in B. No significant pathways were identified for proteins upregulated by mechanomodulation priming. Proteins altered in both low stiffness and oxygen levels were compared and further analyzed (C). The twenty-five common downregulated proteins are involved in ECM organization processes, according to the Reactome database (D; Supplementary Table 9). Abbreviations: Mechanomodulation (Mechano); Benjamini-Hochberg method (BH).

These two proteomics screenings were then directly compared to further understand the similarities between low stiffness and oxygen priming (Figure 4C). Only three proteins were found to be upregulated by both experimental conditions, while six had inverse tendencies (Supplementary Figure 2E). Moreover, the twenty-five proteins identified to be commonly downregulated were described to be involved in ECM organization, corroborating the relevance of ECM for MSCs maintenance under standard culture conditions (Figure 4D and Supplementary Table 9).

### Expanding MSCs under standard culture conditions increases intra and extracellular levels of collagen and other ECM proteins

Next, to determine if the differences observed in the secretory profile of physiologically primed UC-MSCs were due to alterations in the cellular proteome (21), both screenings were compared separately for each experimental condition (Figure 5A-B and C-D, respectively). Priming UC-MSCs on soft platforms decreased the cellular levels of seventeen proteins, which was also reflected in secretome levels. These proteins were found to be involved in ECM organization and signal transduction (Figure 5B and Supplementary Table 10). Moreover, Zyxin and Laminin subunit gamma-1 appear to accumulate within the cell, while Apolipoprotein B-100 is highly secreted (Supplementary Figure 3A-B). Regarding the priming of UC-MSCs at physioxia environments, the results indicate that only three proteins present lower levels in both the intracellular and extracellular space: Fascin, Versican core protein, and Thrombospondin-2, proteins that are recognized to play a role on cell-matrix interactions (Figure 5D). Therefore, it can be concluded that UC-MSCs initiate the readaptation to both physiological environments by modulating the extracellular space.

**Figure 5.**
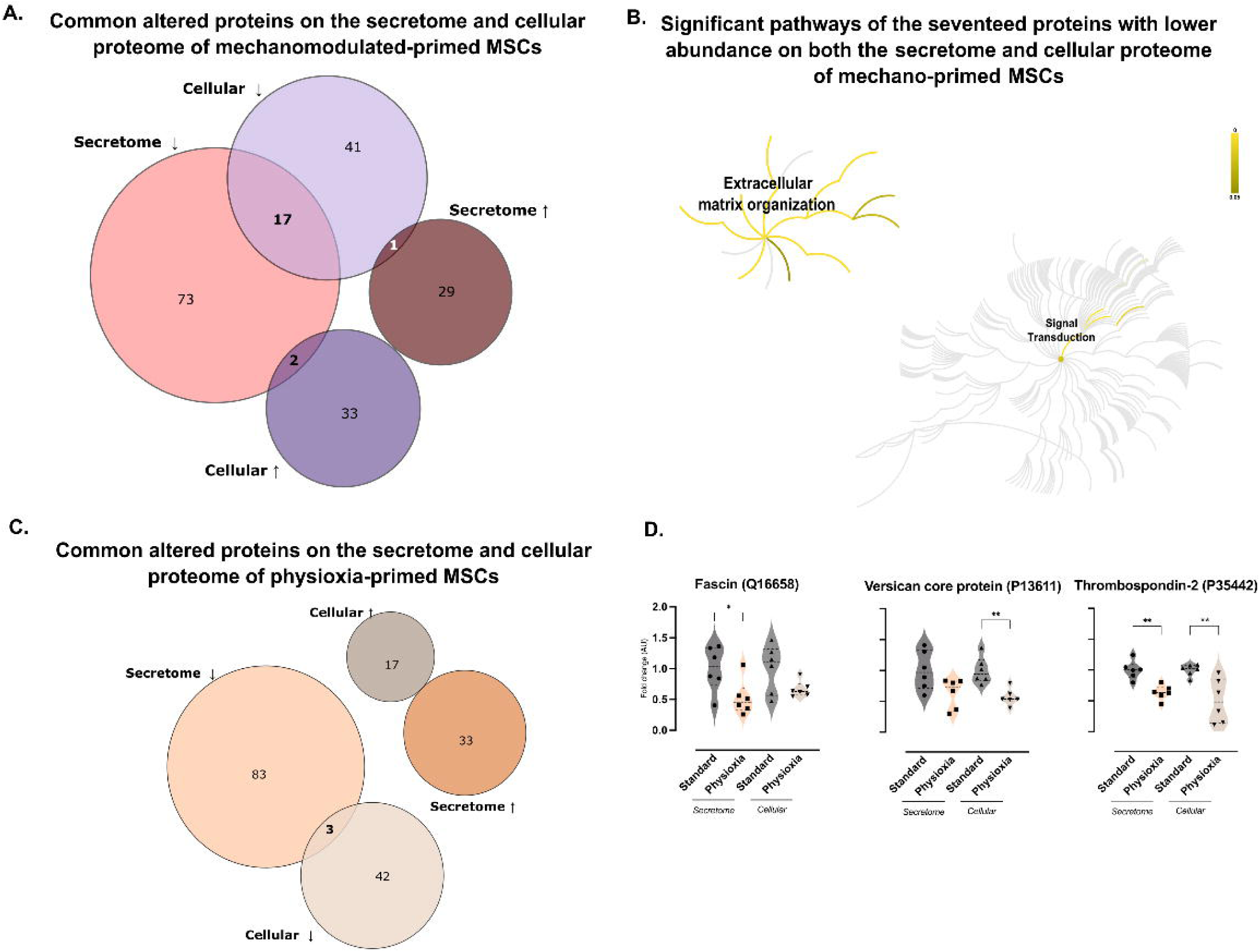
Intra and extracellular levels of ECM proteins decrease when MSCs initiate their readaptation to physiological environments. For each primed method, proteins of interest previously selected for MSCs, cellular proteome, and secretome were compared. Venn diagrams represent common proteins shared by each cellular and secretome fraction for mechanomodulation and physioxia experimental conditions separately (A and C, respectively). Intersections were further explored. The seventeen proteins commonly downregulated proteins upon mechano-priming were analyzed using Reactome to identify significantly altered pathways (yellow) (B, Supplementary Table 10). Regarding physioxia-priming, three proteins were found to be decreased in both intracellular and extracellular spaces (F): Fascin (left), Versican core protein (middle) and Thrombospondin-2 (right) (Mann–Whitney; *p<0.05, **p<0.01).

Finally, to explore if the initial triggered pathways by the priming were kept during extended periods of cell culture, both time points were compared separately for each experimental condition (Figure 6A-B and C-E, respectively). Concerning the mechanical impact on UC-MSCs secretome, short periods increase the secretion of Peroxiredonin-5 (mitochondrial) and N-acetylglucosamine-6-sulfatase, but only the latter protein’s levels are maintained for extended periods (Supplementary Figure 3C, top panel). In contrast, priming decreases the secretion of Endoplasmic reticulum chaperone BIP and Calumenin, whose levels are increased when UC-MSCs are readapted to physiological conditions (Supplementary Figure 3C, bottom panel). However, it appears that the rearrangement at the ECM is maintained over time, as well as PDGF and MET signaling (Figure 6B, Supplementary Table 11). Concerning the impact of time on physioxia-cultured UC-MSCs, cadherin-11 levels increased on physioxia-primed MSCs’ secretome but decreased on readapted MSCs (Supplementary Figure 3D, top panel). Inversely, Tubulin alpha-1A chain, pappalysin-1, and carboxypeptidase A4 levels increased on the secretome of readapted to physioxia MSCs’, but their protein levels were kept low during priming, compared to standard culture conditions, suggestive of a long-term regulation of the secretory profile of these proteins (Supplementary Figure 3D, bottom panel). Nevertheless, the upregulation of proteins associated with the immune system, metabolism of proteins, vesicle-mediated transport, and transport of small molecules pathways seem to be constant at both time points (Figure 6D, Supplementary Table 12), as well as the downregulation of ECM proteins (Figure 6E, Supplementary Table 13). Four of these proteins were shared by primed and UC-MSCs in response to adaptation for low stiffness and oxygen levels, including collagen (Figure 6F). In conclusion, the results suggest that ECM proteins’ levels decrease not only as a response to the initial readaptation of UC-MSCs to a physiological environment but are kept low for long periods, suggesting that they might be essential for maintaining the properties of these cells when expanded under low oxygen or stiffness environments.

**Figure 6.**
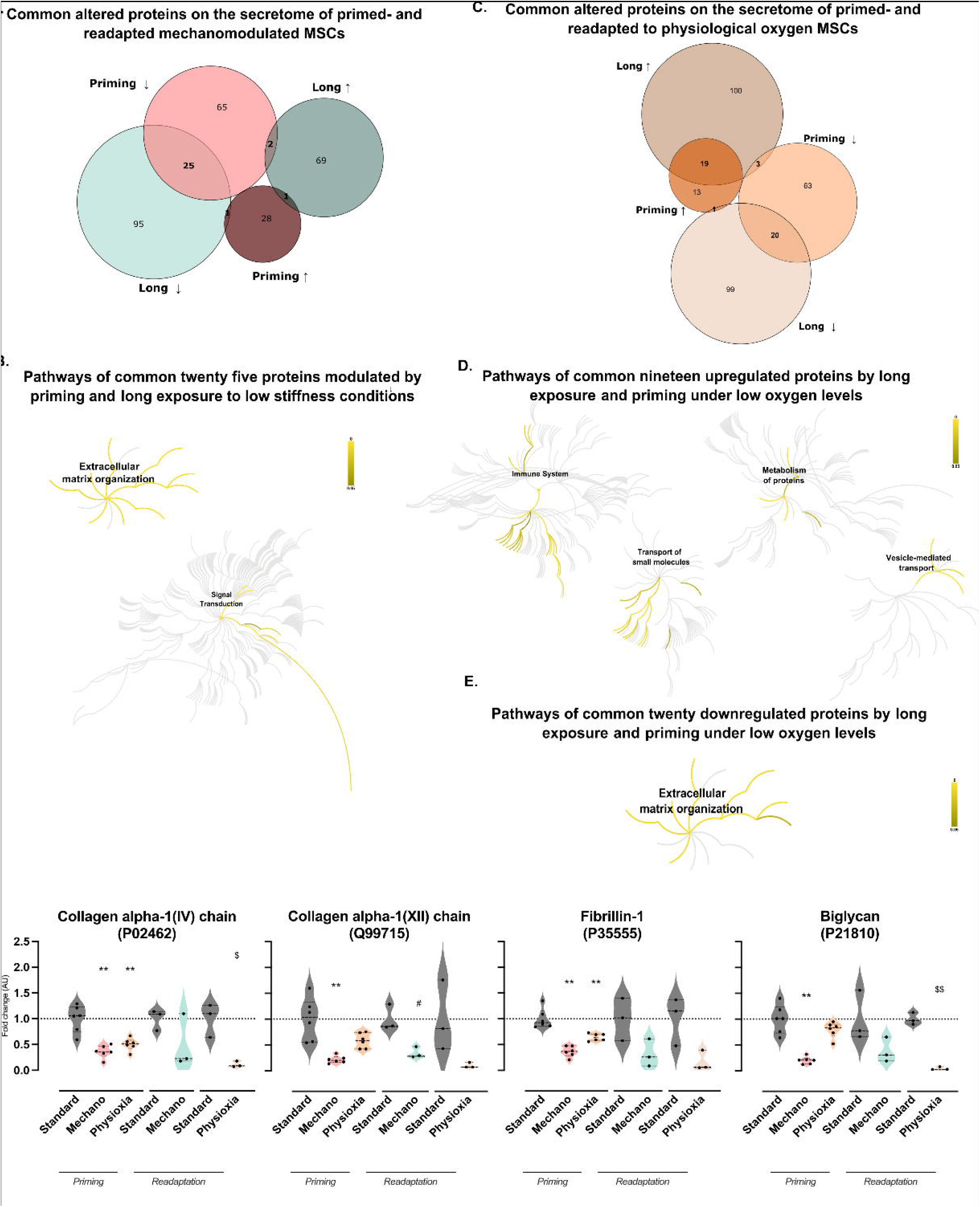
Priming UC-MSCs initiates processes that are kept by readapted MSCs to physiological culture conditions. Statistically different proteins selected for each experimental setup (mechanomodulation and physioxia for primed (48h) and readapted (7-10 days)) were compared (A and C, respectively). Intersections were further explored. The twenty-five proteins that have been downregulated in both periods have significant Reactome pathways (marked in yellow) (B, Supplementary Table 11). Concerning physioxia, the Reactome pathways associated with the common twenty up- and nineteen downregulated proteins are highlighted in yellow (D and E, respectively; Supplementary Tables 12 and 13). Common ECM proteins whose levels were decreased in the secretome by low stiffness or oxygen environments for both primed and readapted UC-MSCs are represented (except biglycan, for physioxia priming, which was not significant) (Mann-Whitney for priming experiments, comparing to the standard:*p<0.05, **p<0.01; t-test for readapted UC-MSCs, comparing to the respective standard: ^#^p<0.05 (for mechanomodulation); ^$^p<0.05, ^$$^p<0.01 (for physioxia).

## Discussion

MSCs exhibit unique properties that allow them to modulate their surrounding environment by secreting molecules with immunomodulatory, antiapoptotic, or antioxidant properties (8, 10). Given the secretome’s well-established beneficial characteristics and ready-to-use availability, there has been a focus on improving and modulating its composition. Published data suggests that reducing the stiffness and oxygen in the culture conditions of UC-MSCs to levels closer to those found in Wharton’s jelly led to rearrangements in protein metabolism and translation processes (21). So, it has been hypothesized that the physiological cues could also alter the secretory profile of these cells.

The proteomic characterization of the CM of UC-MSCs cultured at 3kPa revealed differential secretion of several proteins (Figure 1A and B). Analysis of the biological processes associated with altered proteins indicated an upregulation of several protein disulfide isomerases (PDI) in the secretome (Supplementary Table 14). PDIs play a role in the redox folding within the endoplasmic reticulum (ER), ensuring the formation of disulfide bonds and, consequently, conformational changes in substrates (34). Notably, increased PDI expression has been associated with promoting cell survival in hindlimb ischemia damage, suggesting a putative therapeutic advantage (35). This finding and the higher secretion of antioxidant proteins may be pivotal in modulating the harsh environment typically encountered by UC-MSCs at the injury site (Figure 1C, F). Furthermore, the identification of proteins involved in the TCA cycle and other mitochondrial-specific proteins suggests a possible increase in mitochondrial transfer, a process recognized for its importance in restoring oxidative phosphorylation (36, 37). Mitochondrial activity has previously been shown described to be influenced by matrix stiffness and physioxia, playing a crucial role in the differentiation processes of MSCs by balancing ROS levels and controlling gene expression (38–41). Therefore, future studies should involve a more comprehensive characterization of mitochondria proteins present in the secretome, encompassing their activity and morphology.

Additionally, a decrease in several 40s ribosomal subunits, known for their involvement in translation and rRNA processing mechanisms, was observed in the secretome (Figure 1C, E). The presence of ribosomes in the extracellular environment has already been documented for other cell types (42). The role of ribosomal proteins and RNA composition in regulating gene expression has gained attention in the past few years (43, 44). Hence, exploring the differential secretion of these structures by UC-MSCs and mitochondrial proteins could be a promising approach to investigate the potential implications on the cell signaling mechanisms triggered at the lesion site.

Moreover, culturing UC-MSCs at physioxia deeply altered the secretory profile of these cells for both long and short periods (Figure 2A, B, and Figure 4A). Proteins associated with immunomodulatory processes (both innate and adaptative), particularly immunoglobulins and fibrinolysis, were found to be increased in the CM (Supplementary Tables 3, 5, and 14). In fact, previous studies have shown that exposing MSCs to low levels of oxygen enhances their immunomodulatory properties (45). Conversely, proteins associated with cell adhesion, angiogenesis, or ossification were found to be downregulated. Notably, many of these proteins are ECM components, including collagen, fibronectin, and laminin (Supplementary Table 15). These proteins play a role in determining stem cell fate, suggesting that standard culture conditions may upregulate their secretion and possibly bias their naïve therapeutical properties (46).

Reduced stiffness and oxygen levels decreased ECM protein secretion (Figure 3B and 5B,D). So, to further investigate the commonly modulated secreted proteins, both screenings were compared (Figure 6A,C). The results revealed that collagen was the major component altered (Figure 3B, Figure 6B, E, F; Supplementary Table 16). Collagen fibril crosslinking, which depends on posttranslational modifications, influences tissue tensile strength (47). Higher collagen composition is typically associated with increased tissue elastic modulus, potentially leading to an osteogenic commitment of MSCs (48). Hence, it is intriguing to note that MSCs cultured under physioxia conditions can possibly modulate the stiffness of their microenvironment, which has implications for their cell fate. However, further studies are necessary to analyze the 3D organization of the secreted collagen fibrils since peroxidasin homolog, involved in collagen crosslinking, was also found to be decreased (49). Other matrix constituents, such as fribrilin-1 (P35555) and biglycan (P21810) (Figure 6F), known to determine stem cell fate, were also downregulated (50, 51). In fact, the results suggest that the reorganization of the extracellular space is a key factor in the adaptation of MSCs to standard culture conditions since the differential expression of these components is observed upon priming (48h), particularly in response to mechanical cues (Figure 4B, 5B). Therefore, future work should focus on studying the mechanisms by which alterations in ECM composition affect signaling pathways and their impact on MSC differentiation and therapeutic efficacy. The remodeling of ECM proteins can significantly impact receptor tyrosine kinase signaling pathways, including those mediated by PDGF (Supplementary Tables 10 and 12). Collagen has been shown to target Discoidin domain receptor 1, a collagen-activated receptor tyrosine kinase, thereby modulating cell morphology, ERK activation, and osteogenic commitment (52). Other works have also described the PDGF pathway as being crucial for MSCs’ growth and proliferation (53, 54). Specifically, by interacting with integrins, ECM components can interfere with the PI3K/AKT pathway and alter stem cell fate (Figure 3F) (55). Nevertheless, future research should validate how ECM rearrangements differentially activate these pathways.

The changes in the CM composition induced by the modulation of MSCs’ culture conditions may influence the regenerative potential of the secretome, depending on the target tissue. For instance, a decrease in the levels of ECM proteins essential to promote bone regeneration, such as proteoglycans or C-type lectin (Supplementary Tables 1, 4, and 14), might make the application of physiologically modulated secretome more suitable to other organ systems rather than the skeletal system (56, 57). In addition to common features, a subset of proteins linked to the calnexin/calreticulin cycle (part of the ER chaperon system) were found to be inversely modulated by low stiffness and oxygen levels. These proteins include calreticulin, neutral alpha-glucosidase AB, and protein disulfide-isomerase A3 (Supplementary Figure 1G, Supplementary Table 17). Calreticulin plays a crucial role in mediating osteogenesis, cell adhesion, and fibrotic processes by stimulating fibronectin and collagen deposition in other cell types (58–61). However, the present study suggests that other pathways might also be interfering with ECM deposition since several collagen fibrils were found to be decreased on soft substrates.

In conclusion, analysis of the CM revealed that expanding UC-MSCs under high stiffness and hyperoxia environments (typical of standard culture conditions) may compromise the secretion of PDIs and mitochondrial proteins, as well as proteins involved in immune responses, key players for cell survival at the site of injury. In addition, replicating Wharton’s Jelly stiffness and oxygen levels led to the downregulation of several ECM components. This suggests that maintaining UC-MSCs in an artificial environment might be upregulating these components, biasing stem cell fate, and potentially affecting the therapeutical properties of the secretome.

## List of Abbreviations

CM: Conditioned medium
DDA: Data Dependent Acquisition
DIA: Data Independent Acquisition
ECM: Extracellular matrix
ER: Endoplasmic reticulum
FDR: False Discovery Rate
GO: Gene Ontology
MET: Hepatocyte growth factor receptor
MSCs: Mesenchymal stem cells
PDGF: Platelet-Derived Growth Factor
PDI: Protein disulfide isomerase
PDMS: Polydimethylsiloxane
PLS-DA: Partial Least-Squares Discriminant Analysis
ROS: Reactive oxygen species
SWATH: Sequential Window Acquisition of all Theoretical Mass Spectra
TCA: Tricarboxylic acid cycle
UC: Umbilical cord
UC-MSCs: Umbilical cord mesenchymal stem cells

---

**Supplementary Figure 1.**
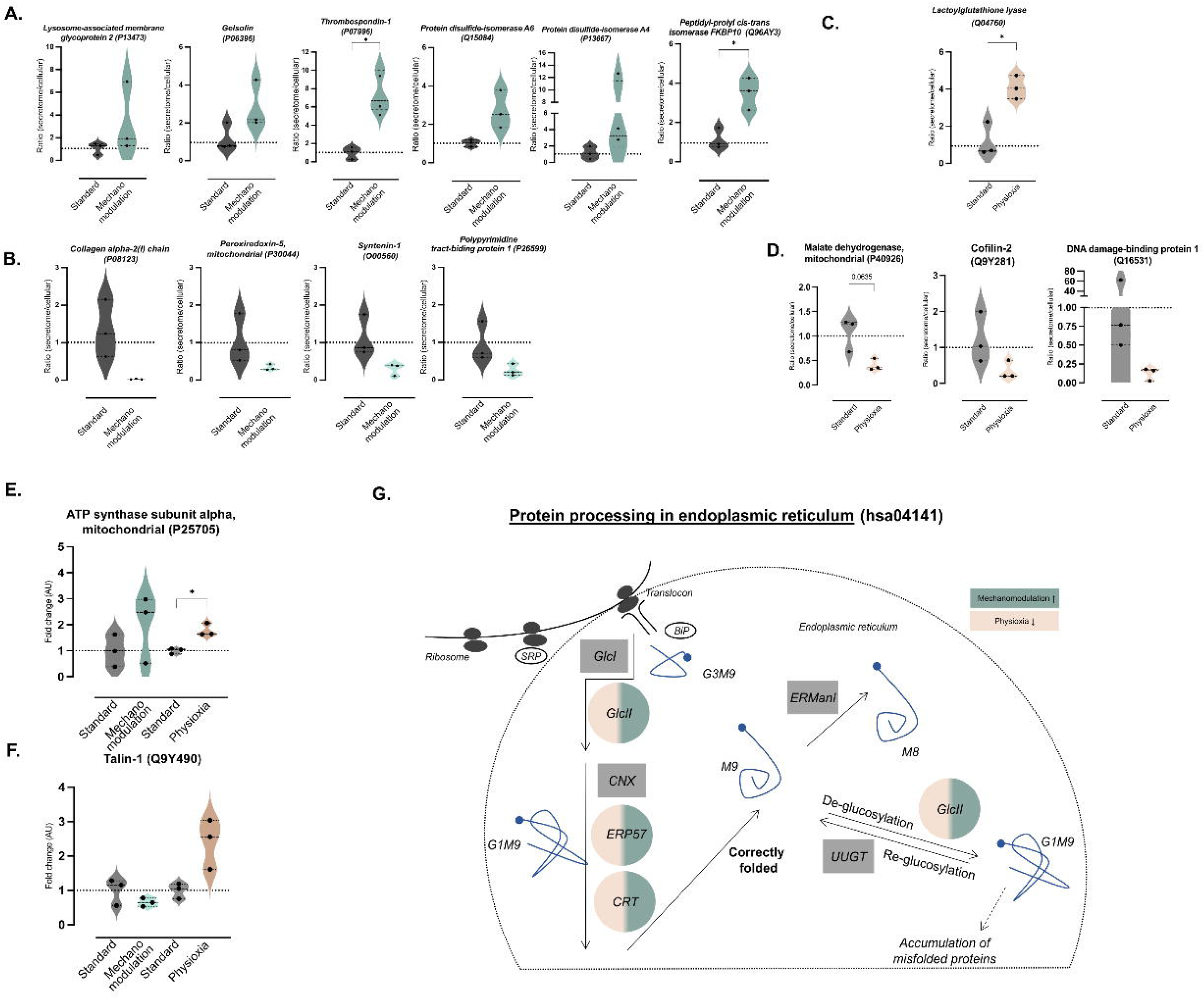
Readaptation of UC-MSCs to physiological conditions differentially modulate intra and extracellular levels of the proteins. For mechanomodulated UC-MSCs, six proteins increased in the secretome but with lower levels on the cellular fraction (A), while four proteins were less secreted but increased levels on the cellular fraction (B) (t-test; *p<0.05). Considering physioxia-adapted UC-MSCs, the protein whose levels are higher in the secretome compared to the cellular proteome, or the three proteins that appear to accumulate within the cells are represented in the Violin plots (C and D, respectively) (t-test; *p<0.05). Comparing both experimental conditions, it was possible to identify ATP synthase subunit alpha (E), whose levels are increased in both systems and Talin-1 (F), which is increased in physioxia and decreased by low stiffness conditions (t-test, *p<0.05). According to the Kegg Pathway database, proteins with opposite tendencies in physiological environments might be involved in the calnexin/calreticulin cycle (G). In detail, the mapped proteins are highlighted in circles (grey: proteins that belong to the pathway; dark green: upregulated on mechanomodulation; light brown: downregulated on physioxia). Abbreviations: SRP: signal recognition particle; BiP: Binding immunoglobulin protein; GlcI: mannosyl-oligosaccharide glucosidase; GlcII: glucosidase II alpha subunit; CNX: calnexin; ERP57: protein disulfide-isomerase A3; CRT: Calreticulin; ERManI: mannosidase alpha class 1A member 2; UUGT: UDP-glucose glycoprotein glucosyltransferase 2.

**Supplementary Figure 2.**
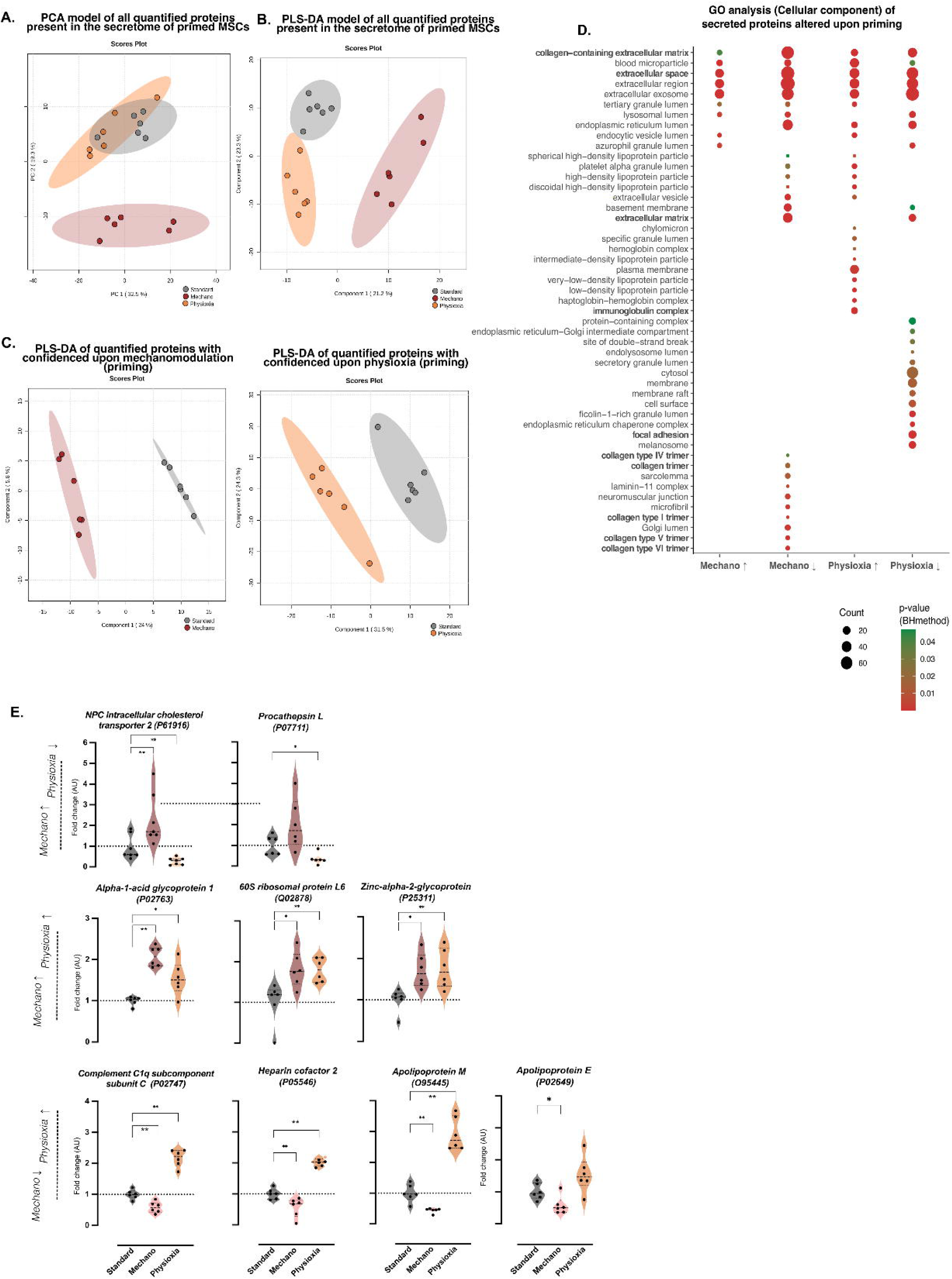
Composition of MSCs secretome is differentially altered by mechanomodulation or physioxia priming. A PCA and a PLS-DA model were generated using all 413 quantified proteins in the three experimental groups (standard (grey), mechano (orange), physioxia (red)) (A, B respectively). Also, the PLS-DA scores plot illustrates mechanomodulation and physioxia priming individually, compared to the standard condition (C, left and right, respectively). Cellular components (GO) of significant proteins (VIP>1, p<0.05, or both; |Log1.5FC|>1) are represented in D. Violin plots represent proteins with inverse tendencies on both experimental conditions (top and bottom) and common upregulated proteins (middle) (E). (Kruskal-Wallis test; *p<0.05, **p<0.01)

**Supplementary Figure 3.**
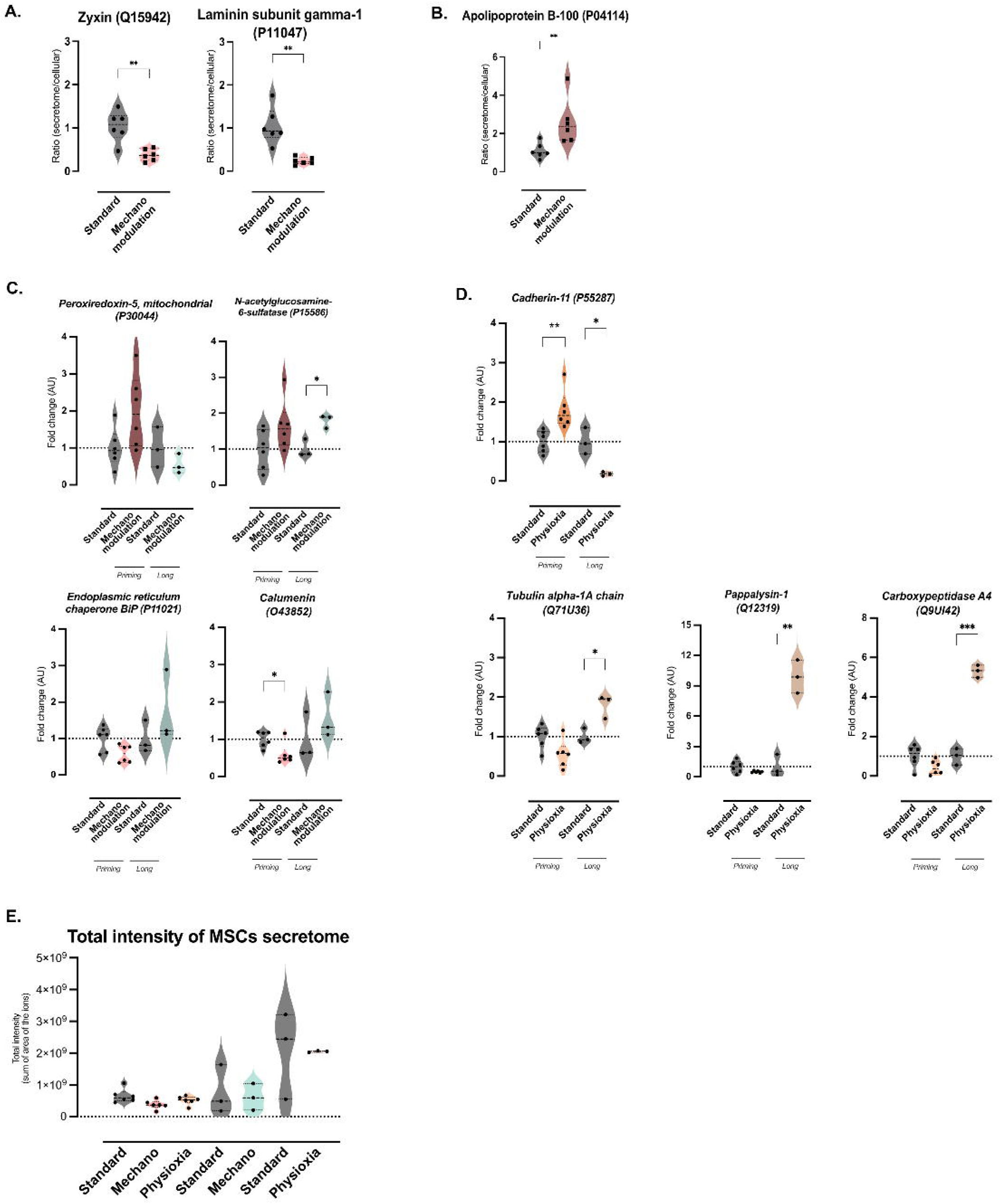
The initial secretory pattern of UC-MSCs is altered during their readaptation to physiological environments. In response to mechanomodulation priming, two proteins appear to accumulate in the intracellular space (A) and the one that appears to be translocated to the extracellular space in MSCs primed by mechanomodulation (B) (Mann–Whitney test; **p<0.01). Primed and readapted UC-MSCs secretome were then compared for each experimental condition. Lowering stiffness for extended periods leads to the upregulation of peroxiredoxin-5 (mitochondrial) and N-acetylglucosamine-6-sulfatase, but the levels of the former were decreased in long-term cultures (C, top panel). Also, two proteins were decreased by mechano-priming, but their levels in the secretome increased for readapted UC-MSCs: endoplasmic reticulum chaperone BIP and calumenin (C, bottom panel) (Mann-Whitney for priming; t-test for readapted MSCs; *p<0.05). Concerning physioxia, Cadherin-11 levels were increased upon priming but decreased after extended periods under physiological oxygen levels (D, top panel). In contrast, three proteins (Tubulin alpha-1A chain, Pappalysin-1 and Carboxypeptidase A4) expression was increased on the long-term experimental setup but decreased on priming (D, bottom panel) (Mann-Whitney for priming; t-test for readapted MSCs; *p<0.05, **p<0.01). The total intensity of the secretome was not altered by the experimental conditions (E).

## Declaration of generative AI and AI-assisted technologies in the writing process

During the preparation of this work the authors used ChatGPT in order to check spelling and correct grammar errors. After using this tool/service, the authors reviewed and edited the content as needed and take full responsibility for the content of the publication.

