## Supplementary Tables for "Readaptation of mesenchymal stem cells to high stiffness and oxygen environments modulate the extracellular matrix"

| Supplementary Table 1 – Significant Reactome pathways of commonly downregulated proteins on the secretome and proteome of mechanomodulated MSCs (readaptation) | | | |
| --- | --- | --- | --- |
| **Reactome Pathway** | **Identifier** | **FDR** | **ID mapped** |
| *Translation initiation complex formation* | R-HSA-72649 | 4.66E-14 | Q04637;P62277;P15880;P46781;P62701;P11940;P46783;P62081;P08865;P60866;P62249 |
| *Activation of the mRNA upon binding of the cap-binding complex and eIFs, and subsequent binding to 43S* | R-HSA-72662 | 4.66E-14 | Q04637;P62277;P15880;P46781;P62701;P11940;P46783;P62081;P08865;P60866;P62249 |
| ***Nonsense Mediated Decay (NMD) independent of the Exon Junction Complex (EJC)*** | R-HSA-975956 | 7.77E-14 | Q04637;P62277;P15880;P61353;P46781;P62701;P11940;P46783;P62081;P08865;P60866;P62249 |
| *L13a-mediated translational silencing of Ceruloplasmin expression* | R-HSA-156827 | 3.08E-13 | Q04637;P62277;P15880;P61353;P46781;P62701;P11940;P46783;P62081;P08865;P60866;P62249 |
| *SARS-CoV-2 modulates host translation machinery* | R-HSA-9754678 | 3.08E-13 | P62277;P15880;P46781;P62701;P46783;P62081;P62318;P08865;P60866;P62249 |
| *Nonsense-Mediated Decay (NMD)* | R-HSA-927802 | 3.73E-13 | Q04637;P62277;P15880;P61353;P46781;P62701;P11940;P46783;P62081;P08865;P60866;P62249 |
| *Nonsense Mediated Decay (NMD) enhanced by the Exon Junction Complex (EJC)* | R-HSA-975957 | 3.73E-13 | Q04637;P62277;P15880;P61353;P46781;P62701;P11940;P46783;P62081;P08865;P60866;P62249 |
| *Cap-dependent Translation Initiation* | R-HSA-72737 | 3.83E-13 | Q04637;P62277;P15880;P61353;P46781;P62701;P11940;P46783;P62081;P08865;P60866;P62249 |
| ***Eukaryotic Translation Initiation*** | R-HSA-72613 | 3.83E-13 | Q04637;P62277;P15880;P61353;P46781;P62701;P11940;P46783;P62081;P08865;P60866;P62249 |
| *Ribosomal scanning and start codon recognition* | R-HSA-72702 | 4.43E-13 | Q04637;P62277;P15880;P46781;P62701;P46783;P62081;P08865;P60866;P62249 |
| *Regulation of expression of SLITs and ROBOs* | R-HSA-9010553 | 7.00E-13 | P46781;P11940;P62081;Q04637;P62277;P15880;P61353;P62701;P46783;P61289;P08865;P60866;P62249 |
| *SARS-CoV-1 modulates host translation machinery* | R-HSA-9735869 | 7.19E-13 | P62277;P15880;P46781;P62701;P46783;P62081;P08865;P60866;P62249 |
| *Axon guidance* | R-HSA-422475 | 3.76E-12 | P02461;P46781;P11940;Q13509;P62081;P12111;P15311;Q04637;P12110;P62277;P15880;P61353;P62701;P46783;P61289;P08865;P60866;P62249 |
| *GTP hydrolysis and joining of the 60S ribosomal subunit* | R-HSA-72706 | 5.05E-12 | Q04637;P62277;P15880;P61353;P46781;P62701;P46783;P62081;P08865;P60866;P62249 |
| *Formation of the ternary complex, and, subsequently, the 43S complex* | R-HSA-72695 | 5.93E-12 | P62277;P15880;P46781;P62701;P46783;P62081;P08865;P60866;P62249 |
| *Nervous system development* | R-HSA-9675108 | 6.59E-12 | P02461;P46781;P11940;Q13509;P62081;P12111;P15311;Q04637;P12110;P62277;P15880;P61353;P62701;P46783;P61289;P08865;P60866;P62249 |
| *Signaling by ROBO receptors* | R-HSA-376176 | 8.66E-12 | P46781;P11940;P62081;Q04637;P62277;P15880;P61353;P62701;P46783;P61289;P08865;P60866;P62249 |
| *Peptide chain elongation* | R-HSA-156902 | 1.53E-11 | P62277;P15880;P61353;P46781;P62701;P46783;P62081;P08865;P60866;P62249 |
| ***Eukaryotic Translation Termination*** | R-HSA-72764 | 2.14E-11 | P62277;P15880;P61353;P46781;P62701;P46783;P62081;P08865;P60866;P62249 |
| *Selenocysteine synthesis* | R-HSA-2408557 | 2.14E-11 | P62277;P15880;P61353;P46781;P62701;P46783;P62081;P08865;P60866;P62249 |
| ***Eukaryotic Translation Elongation*** | R-HSA-156842 | 2.26E-11 | P62277;P15880;P61353;P46781;P62701;P46783;P62081;P08865;P60866;P62249 |
| *Viral mRNA Translation* | R-HSA-192823 | 3.84E-11 | P62277;P15880;P61353;P46781;P62701;P46783;P62081;P08865;P60866;P62249 |
| *Formation of a pool of free 40S subunits* | R-HSA-72689 | 3.84E-11 | P62277;P15880;P61353;P46781;P62701;P46783;P62081;P08865;P60866;P62249 |
| *Response of EIF2AK4 (GCN2) to amino acid deficiency* | R-HSA-9633012 | 3.84E-11 | P62277;P15880;P61353;P46781;P62701;P46783;P62081;P08865;P60866;P62249 |
| *SRP-dependent cotranslational protein targeting to membrane* | R-HSA-1799339 | 9.81E-11 | P62277;P15880;P61353;P46781;P62701;P46783;P62081;P08865;P60866;P62249 |
| *Selenoamino acid metabolism* | R-HSA-2408522 | 1.49E-10 | P62277;P15880;P61353;P46781;P62701;P46783;P62081;P08865;P60866;P62249 |
| *Influenza Viral RNA Transcription and Replication* | R-HSA-168273 | 1.62E-09 | P62277;P15880;P61353;P46781;P62701;P46783;P62081;P08865;P60866;P62249 |
| *SARS-CoV-1-host interactions* | R-HSA-9692914 | 2.07E-09 | P62277;P15880;P46781;P62701;P46783;P62081;P08865;P60866;P62249 |
| *Cellular response to starvation* | R-HSA-9711097 | 2.07E-09 | P62277;P15880;P61353;P46781;P62701;P46783;P62081;P08865;P60866;P62249 |
| ***Translation*** | R-HSA-72766 | 3.63E-09 | Q04637;P62277;P15880;P61353;P46781;P62701;P11940;P46783;P62081;P08865;P60866;P62249 |
| *Influenza Infection* | R-HSA-168255 | 4.61E-09 | P62277;P15880;P61353;P46781;P62701;P46783;P62081;P08865;P60866;P62249 |
| *Major pathway of rRNA processing in the nucleolus and cytosol* | R-HSA-6791226 | 8.34E-09 | P62277;P15880;P61353;P46781;P62701;P46783;P62081;P08865;P60866;P62249 |
| *rRNA processing in the nucleus and cytosol* | R-HSA-8868773 | 1.28E-08 | P62277;P15880;P61353;P46781;P62701;P46783;P62081;P08865;P60866;P62249 |
| ***Metabolism of RNA*** | R-HSA-8953854 | 1.45E-08 | P46781;P11940;P62081;Q04637;P22626;P62277;P15880;P61353;P62701;P46783;P62318;P61289;P61978;P08865;P60866;P62249 |
| ***rRNA processing*** | R-HSA-72312 | 2.07E-08 | P62277;P15880;P61353;P46781;P62701;P46783;P62081;P08865;P60866;P62249 |
| *SARS-CoV-1 Infection* | R-HSA-9678108 | 3.17E-08 | P62277;P15880;P46781;P62701;P46783;P62081;P08865;P60866;P62249 |
| *SARS-CoV-2-host interactions* | R-HSA-9705683 | 4.24E-08 | P62277;P15880;P46781;P62701;P46783;P62081;P62318;P08865;P60866;P62249 |
| *Developmental Biology* | R-HSA-1266738 | 4.48E-07 | P02461;P46781;P11940;Q13509;P62081;P12111;P15311;Q04637;P12110;P62277;P15880;P61353;P62701;P46783;P61289;P08865;P60866;P62249 |
| *Metabolism of amino acids and derivatives* | R-HSA-71291 | 4.97E-07 | P62277;P15880;P61353;P46781;P62701;P46783;P62081;P61289;P08865;P60866;P62249 |
| *SARS-CoV-2 Infection* | R-HSA-9694516 | 1.09E-06 | P62277;P15880;P46781;P62701;P46783;P62081;P62318;P08865;P60866;P62249 |
| *Viral Infection Pathways* | R-HSA-9824446 | 5.48E-06 | P46781;Q13509;P62081;P62277;P13010;P15880;P61353;P62701;P46783;P62318;P61289;P61978;P08865;P60866;P62249 |
| *Assembly of collagen fibrils and other multimeric structures* | R-HSA-2022090 | 2.49E-05 | P02461;Q15149;P12110;Q92626;P12111 |
| *SARS-CoV Infections* | R-HSA-9679506 | 7.17E-05 | P62277;P15880;P46781;P62701;P46783;P62081;P62318;P08865;P60866;P62249 |
| *Cellular responses to stress* | R-HSA-2262752 | 8.83E-05 | P62277;P15880;P61353;P46781;P62701;Q13509;P46783;P62081;P61289;P08865;P60866;P62249 |
| *Cellular responses to stimuli* | R-HSA-8953897 | 1.06E-04 | P62277;P15880;P61353;P46781;P62701;Q13509;P46783;P62081;P61289;P08865;P60866;P62249 |
| *Infectious disease* | R-HSA-5663205 | 1.29E-04 | P46781;Q13509;P62081;P62277;P13010;P15880;P61353;P62701;P46783;P62318;P61289;P61978;P08865;P60866;P62249 |
| ***Collagen formation*** | R-HSA-1474290 | 1.29E-04 | P02461;Q15149;P12110;Q92626;P12111 |
| *Disease* | R-HSA-1643685 | 8.86E-04 | Q8IVT5;P46781;Q13509;P62081;P33176;P62277;P13010;P15880;P61353;P62701;P46783;P62318;Q16543;P61289;P61978;P08865;P60866;P62249 |
| ***ECM proteoglycans*** | R-HSA-3000178 | 1.20E-03 | P02461;P12110;P05121;P12111 |
| ***Metabolism of proteins*** | R-HSA-392499 | 1.36E-03 | P46781;P11940;Q13509;P62081;P33176;Q04637;P62277;P15880;P61353;P62701;P46783;Q15582;P61289;P61978;P08865;P60866;P62249 |
| *NCAM1 interactions* | R-HSA-419037 | 3.58E-03 | P02461;P12110;P12111 |
| *Collagen chain trimerization* | R-HSA-8948216 | 4.10E-03 | P02461;P12110;P12111 |
| ***Extracellular matrix organization*** | R-HSA-1474244 | 4.18E-03 | P02461;Q15149;P12110;Q92626;P05121;P12111 |
| *Z-decay: degradation of maternal mRNAs by zygotically expressed factors* | R-HSA-9820865 | 5.90E-03 | Q04637;P11940 |
| *AUF1 (hnRNP D0) binds and destabilizes mRNA* | R-HSA-450408 | 7.18E-03 | Q04637;P11940;P61289 |
| *rRNA modification in the nucleus and cytosol* | R-HSA-6790901 | 8.74E-03 | P15880;P46781;P62081 |
| *Signaling by PDGF* | R-HSA-186797 | 8.74E-03 | P02461;P12110;P12111 |
| *Collagen degradation* | R-HSA-1442490 | 1.05E-02 | P02461;P12110;P12111 |
| *NCAM signaling for neurite out-growth* | R-HSA-375165 | 1.05E-02 | P02461;P12110;P12111 |
| *Collagen biosynthesis and modifying enzymes* | R-HSA-1650814 | 1.20E-02 | P02461;P12110;P12111 |
| *Integrin cell surface interactions* | R-HSA-216083 | 2.01E-02 | P02461;P12110;P12111 |
| *Deadenylation of mRNA* | R-HSA-429947 | 2.13E-02 | Q04637;P11940 |
| *M-decay: degradation of maternal mRNAs by maternally stored factors* | R-HSA-9820841 | 2.13E-02 | Q04637;P11940 |
| *Regulation of mRNA stability by proteins that bind AU-rich elements* | R-HSA-450531 | 2.21E-02 | Q04637;P11940;P61289 |
| *RHOH GTPase cycle* | R-HSA-9013407 | 4.55E-02 | Q15758;P02786 |

| Supplementary Table 2 – Significant Reactome pathways of commonly upregulated proteins on the secretome and proteome of mechanomodulated MSCs (readaptation) | | | |
| --- | --- | --- | --- |
| **Reactome Pathway** | **Identifier** | **FDR** | **ID mapped** |
| ***Mitochondrial protein import*** | R-HSA-1268020 | 3.72E-04 | O75390;P25705;P10809;P06576 |
| ***Metabolism of carbohydrates*** | R-HSA-71387 | 3.72E-04 | P15121;P00505;Q04446;P40926;P19367;P00352 |
| ***Mitochondrial biogenesis*** | R-HSA-1592230 | 6.25E-04 | Q04837;P25705;P06576;P04179 |
| ***Detoxification of Reactive Oxygen Species*** | R-HSA-3299685 | 1.01E-03 | P30048;P04179;Q16881 |
| *Fructose metabolism* | R-HSA-5652084 | 1.40E-03 | P15121;P00352 |
| *Transcriptional activation of mitochondrial biogenesis* | R-HSA-2151201 | 2.02E-03 | Q04837;P06576;P04179 |
| *Protein localization* | R-HSA-9609507 | 2.13E-03 | O75390;P25705;P10809;P06576 |
| ***The citric acid (TCA) cycle and respiratory electron transport*** | R-HSA-1428517 | 2.44E-03 | O75390;P25705;P40926;P06576 |
| *Formation of ATP by chemiosmotic coupling* | R-HSA-163210 | 5.19E-03 | P25705;P06576 |
| *Glucose metabolism* | R-HSA-70326 | 5.94E-03 | P00505;P40926;P19367 |
| *Citric acid cycle (TCA cycle)* | R-HSA-71403 | 5.94E-03 | O75390;P40926 |
| ***Metabolism*** | R-HSA-1430728 | 7.24E-03 | O75390;P15121;P25705;P00505;Q04446;P40926;P19367;P00352;P06576;Q16881 |
| *Cristae formation* | R-HSA-8949613 | 9.55E-03 | P25705;P06576 |
| *Organelle biogenesis and maintenance* | R-HSA-1852241 | 9.55E-03 | Q04837;P25705;P06576;P04179 |
| *Gluconeogenesis* | R-HSA-70263 | 9.75E-03 | P00505;P40926 |
| *Metabolism of ingested MeSeO_2_H into MeSeH* | R-HSA-5263617 | 9.75E-03 | Q16881 |
| *Defective HK1 causes hexokinase deficiency (HK deficiency)* | R-HSA-5619056 | 9.75E-03 | P19367 |
| *Fructose biosynthesis* | R-HSA-5652227 | 1.95E-02 | P15121 |
| ***Pyruvate metabolism and Citric Acid (TCA) cycle*** | R-HSA-71406 | 2.16E-02 | O75390;P40926 |
| *Glycogen storage disease type IV (GBE1)* | R-HSA-3878781 | 2.43E-02 | Q04446 |
| ***Cellular response to chemical stress*** | R-HSA-9711123 | 2.46E-02 | P30048;P04179;Q16881 |
| *Metabolism of ingested H_2_SeO_4_ and H_2_SeO_3_ into H_2_Se* | R-HSA-2408550 | 3.24E-02 | Q16881 |
| *Fructose catabolism* | R-HSA-70350 | 3.24E-02 | P00352 |
| *TFAP2A acts as a transcriptional repressor during retinoic acid induced cell differentiation* | R-HSA-8869496 | 3.24E-02 | P10809 |
| *Uptake and function of diphtheria toxin* | R-HSA-5336415 | 4.53E-02 | Q16881 |

| *Supplementary Table 3 – Significant Biological processes of upregulated secretome proteins by physioxia (long exposure)* | | | |
| --- | --- | --- | --- |
| **Biological process** | **Fold Enrichment** | **BH method** | **IDs mapped** |
| *complement activation, classical pathway* | 40.21 | 3.92E-38 | A0A0B4J1Y9; P01877; P04003; P02747; P02746; P01859; P01871; A0A0C4DH38; P01860; P01857; P01861; P01834; P0C0L5; B9A064; P02748; P01031; P01880; P06331; P05155; P01876; A0A0C4DH36; P01743; P02743; P0C0L4; P07357; A0A0B4J1X5; A0A0B4J1V6; A0A0B4J1V0; P0DP03; A0A075B6Q5; P01024; |
| *phagocytosis, engulfment* | 29.42 | 5.38E-22 | A0A0B4J1Y9; P01877; P01859; P01871; A0A0C4DH38; P01860; P01857; P01861; P01834; B9A064; P01880; P06331; P01876; A0A0C4DH36; P01743; A0A0B4J1X5; A0A0B4J1V6; A0A0B4J1V0; P35579; P0DP03; A0A075B6Q5; |
| *phagocytosis, recognition* | 33.36 | 5.38E-22 | A0A0B4J1Y9; P01877; P01859; P01871; A0A0C4DH38; P01860; P01857; P01861; P01834; B9A064; P01880; P06331; P01876; A0A0C4DH36; P01743; A0A0B4J1X5; A0A0B4J1V6; A0A0B4J1V0; P0DP03; A0A075B6Q5; |
| *positive regulation of B cell activation* | 34.17 | 5.38E-22 | A0A0B4J1Y9; P01877; P01859; P01871; A0A0C4DH38; P01860; P01857; P01861; P01834; B9A064; P01880; P06331; P01876; A0A0C4DH36; P01743; A0A0B4J1X5; A0A0B4J1V6; A0A0B4J1V0; P0DP03; A0A075B6Q5; |
| *B cell receptor signaling pathway* | 24.80 | 2.58E-19 | A0A0B4J1Y9; P01877; P01859; P01871; A0A0C4DH38; P01860; P01857; P01861; P01834; B9A064; P01880; P06331; P01876; A0A0C4DH36; P01743; A0A0B4J1X5; A0A0B4J1V6; A0A0B4J1V0; P0DP03; A0A075B6Q5; |
| *innate immune response* | 7.91 | 2.54E-16 | A0A0B4J1Y9; P01877; P04003; P02747; P02746; P01859; P01871; A0A0C4DH38; P18428; P01860; P01857; P01861; P01834; P0C0L5; B9A064; P01880; P06331; P05155; P01876; A0A0C4DH36; P01743; P00748; P02743; P0C0L4; P04083; A0A0B4J1X5; A0A0B4J1V6; A0A0B4J1V0; P0DP03; A0A075B6Q5; O14791; |
| *defense response to bacterium* | 14.86 | 9.56E-16 | A0A0B4J1Y9; P01877; P01859; P01871; A0A0C4DH38; P01860; P01857; P00738; P01861; P01834; B9A064; P01880; P06331; P01876; A0A0C4DH36; P01743; A0A0B4J1X5; A0A0B4J1V6; A0A0B4J1V0; P0DP03; A0A075B6Q5; |
| *negative regulation of endopeptidase activity* | 32.34 | 2.50E-12 | P02765; P01042; P05543; P01009; P29622; P05546; P04196; P04004; P01011; P01008; P01019; P05155; |
| *blood coagulation* | 23.06 | 1.41E-11 | P00740; P00734; P00742; P01042; P01009; P05546; P01008; P05155; P00748; P00747; P08758; P07225; P03952; |
| *immune response* | 7.94 | 1.41E-11 | P01877; P02747; A0A0C4DH72; A0A075B6H7; P04004; A0A0C4DH25; P01701; P01834; P80748; P01880; P06331; P01876; P01743; P0DP09; P01602; P06312; P01611; P07357; A0A075B6R9; P01703; A0A087WSY6; P04433; P01024; |
| *fibrinolysis* | 59.01 | 4.23E-10 | P00734; P04196; P07355; P05155; P00748; P00747; P07225; P03952; |
| *acute-phase response* | 32.34 | 6.47E-09 | P02765; P02763; Q14624; P00734; P01009; P18428; P00738; P01011; P02743; |
| *zymogen activation* | 40.06 | 5.51E-06 | P00740; O43866; P00739; P00738; P00748; P03952; |
| *retina homeostasis* | 23.94 | 1.28E-05 | P01877; P02787; P01860; P01834; P01876; P02768; P60709; |
| *adaptive immune response* | 5.30 | 4.55E-05 | A0A0C4DH72; A0A075B6H7; P01871; A0A0C4DH25; P01701; P80748; P0DP09; P01602; P06312; P01611; P04083; A0A075B6R9; P01703; A0A087WSY6; P0DSN7; P04433; |
| *complement activation* | 41.25 | 7.60E-05 | P02746; P0C0L5; P0C0L4; P07357; P01024; |
| *positive regulation of cholesterol esterification* | 62.33 | 2.20E-04 | P06727; P02647; P01019; P02649; |
| *negative regulation of fibrinolysis* | 56.10 | 3.44E-04 | P00734; P04196; P02749; P00747; |
| *cholesterol efflux* | 28.05 | 5.25E-04 | P06727; O95445; P02647; P02649; P04114; |
| *platelet aggregation* | 18.29 | 5.34E-04 | Q13418; Q86UX7; P68871; Q9Y490; P60709; P35579; |
| *high-density lipoprotein particle assembly* | 46.76 | 6.87E-04 | P06727; O95445; P02647; P02649; |
| *vitamin transport* | 105.12 | 7.26E-04 | P02647; P02774; P43652; |
| *lipoprotein biosynthetic process* | 105.12 | 7.26E-04 | P02647; P02649; P04114; |
| *positive regulation of fibrinolysis* | 105.12 | 7.26E-04 | P00748; P00747; P03952; |
| *negative regulation of blood coagulation* | 43.16 | 8.29E-04 | P01042; P02749; P04004; P02649; |
| *complement activation, alternative pathway* | 37.41 | 1.51E-03 | P02748; P01031; P07357; P01024; |
| *high-density lipoprotein particle remodeling* | 35.08 | 1.92E-03 | P06727; O95445; P02647; P02649; |
| *reverse cholesterol transport* | 33.01 | 2.41E-03 | P06727; O95445; P02647; P02649; |
| *positive regulation of apoptotic cell clearance* | 70.14 | 2.97E-03 | P0C0L5; P0C0L4; P01024; |
| *high-density lipoprotein particle clearance* | 60.13 | 5.00E-03 | O95445; P02647; P02649; |
| *lipoprotein metabolic process* | 26.73 | 5.35E-03 | P06727; O95445; P02647; O14791; |
| *cholesterol metabolic process* | 10.79 | 7.60E-03 | P06727; P02647; P27169; P02649; P04114; O14791; |
| *positive regulation of lipoprotein lipase activity* | 46.78 | 1.05E-02 | P06727; P02647; P02749; |
| *very-low-density lipoprotein particle remodeling* | 46.78 | 1.05E-02 | P06727; P02647; P02649; |
| *negative regulation of complement activation, lectin pathway* | 140.04 | 1.64E-02 | P01023; P05155; |
| *negative regulation of plasma lipoprotein oxidation* | 140.04 | 1.64E-02 | P06727; O95445; |
| *detection of molecule of bacterial origin* | 140.04 | 1.64E-02 | P18428; P0C0L5; |
| *Factor XII activation* | 140.04 | 1.64E-02 | P00748; P03952; |
| *hyaluronan metabolic process* | 38.28 | 1.78E-02 | Q14624; Q06033; P19827; |
| *phospholipid efflux* | 35.10 | 2.23E-02 | P06727; P02647; P02649; |
| *negative regulation of smooth muscle cell proliferation* | 17.54 | 2.23E-02 | Q13418; P17936; P05090; P02649; |
| *cytolysis by host of symbiont cells* | 35.10 | 2.23E-02 | P00734; P04196; O14791; |
| *blood coagulation, intrinsic pathway* | 93.51 | 4.25E-02 | P02749; P00748; |
| *positive regulation of blood coagulation* | 28.08 | 4.25E-02 | P00734; P02749; P00748; |

| Supplementary Table 4 – Significant Reactome pathways of commonly downregulated proteins on the secretome and proteome of MSCs cultured under physioxia (long exposure) | | | |
| --- | --- | --- | --- |
| **Reactome Pathway** | **Identifier** | **FDR** | **ID mapped** |
| ***ECM proteoglycans*** | R-HSA-3000178 | 3.40E-09 | P02452 (Collagen alpha-1(I) chain);P02461(Collagen alpha-1(III) chain);P08123 (Collagen alpha-2(I) chain);P20908 (Collagen alpha-1(V) chain);P13611 (Versican core protein);P51884 (Lumican);P02751 (Fibronectin);P11047(Laminin subunit gamma-1) |
| ***Extracellular matrix organization*** | R-HSA-1474244 | 1.10E-08 | P02452;Q9Y6C2;P02461;P08123;P35555;P20908;P08253;P13611;P51884;P11047;P02751 |
| *MET activates PTK2 signaling* | R-HSA-8874081 | 1.10E-08 | P02452;P02461;P08123;P20908;P02751;P11047 |
| *MET promotes cell motility* | R-HSA-8875878 | 5.27E-08 | P02452;P02461;P08123;P20908;P02751;P11047 |
| *Integrin cell surface interactions* | R-HSA-216083 | 6.71E-08 | P02452;P02461;P08123;P35555;P20908;P51884;P02751 |
| *Degradation of the extracellular matrix* | R-HSA-1474228 | 6.71E-08 | P02452;P02461;P08123;P35555;P20908;P08253;P02751;P11047 |
| *Non-integrin membrane-ECM interactions* | R-HSA-3000171 | 2.24E-07 | P02452;P02461;P08123;P20908;P02751;P11047 |
| *Syndecan interactions* | R-HSA-3000170 | 2.24E-07 | P02452;P02461;P08123;P20908;P02751 |
| *Signaling by MET* | R-HSA-6806834 | 1.17E-06 | P02452;P02461;P08123;P20908;P02751;P11047 |
| *Scavenging by Class A Receptors* | R-HSA-3000480 | 3.71E-06 | P02452;P02461;P08123;P27797 |
| *Collagen degradation* | R-HSA-1442490 | 1.13E-05 | P02452;P02461;P08123;P20908;P08253 |
| *Regulation of Insulin-like Growth Factor (IGF) transport and uptake by Insulin-like Growth Factor Binding Proteins (IGFBPs)* | R-HSA-381426 | 1.16E-05 | P09382;P35555;P13611;P08253;P02751;P11047 |
| *Collagen chain trimerization* | R-HSA-8948216 | 7.66E-05 | P02452;P02461;P08123;P20908 |
| ***Post-translational protein phosphorylation*** | R-HSA-8957275 | 1.06E-04 | P09382;P35555;P13611;P02751;P11047 |
| *Assembly of collagen fibrils and other multimeric structures* | R-HSA-2022090 | 2.38E-04 | P02452;P02461;P08123;P20908 |
| ***Collagen biosynthesis and modifying enzymes*** | R-HSA-1650814 | 3.16E-04 | P02452;P02461;P08123;P20908 |
| ***Calnexin/calreticulin cycle*** | R-HSA-901042 | 5.14E-04 | Q14697;P30101;P27797 |
| ***Collagen formation*** | R-HSA-1474290 | 9.06E-04 | P02452;P02461;P08123;P20908 |
| *N-glycan trimming in the ER and Calnexin/Calreticulin cycle* | R-HSA-532668 | 1.03E-03 | Q14697;P30101;P27797 |
| *Molecules associated with elastic fibres* | R-HSA-2129379 | 1.21E-03 | Q9Y6C2;P35555;P02751 |
| *Platelet Aggregation (Plug Formation)* | R-HSA-76009 | 1.37E-03 | P02452;P08123;P02751 |
| *Diseases associated with glycosaminoglycan metabolism* | R-HSA-3560782 | 1.47E-03 | P13611;P07686;P51884 |
| ***Elastic fibre formation*** | R-HSA-1566948 | 1.61E-03 | Q9Y6C2;P35555;P02751 |
| *Binding and Uptake of Ligands by Scavenger Receptors* | R-HSA-2173782 | 2.59E-03 | P02452;P02461;P08123;P27797 |
| *GP1b-IX-V activation signalling* | R-HSA-430116 | 3.60E-03 | P02452;P08123 |
| *Platelet activation, signaling and aggregation* | R-HSA-76002 | 3.74E-03 | P02452;Q08380;P08123;P02751;P10909 |
| *CS/DS degradation* | R-HSA-2024101 | 4.20E-03 | P13611;P07686 |
| *Platelet Adhesion to exposed collagen* | R-HSA-75892 | 4.20E-03 | P02452;P08123 |
| *Keratan sulfate degradation* | R-HSA-2022857 | 4.20E-03 | P07686;P51884 |
| *Anchoring fibril formation* | R-HSA-2214320 | 4.20E-03 | P02452;P08123 |
| *Crosslinking of collagen fibrils* | R-HSA-2243919 | 6.02E-03 | P02452;P08123 |
| ***Hemostasis*** | R-HSA-109582 | 1.23E-02 | P02452;P07093;Q08380;P14174;P08123;P02751;P10909 |
| ***Signaling by Receptor Tyrosine Kinases*** | R-HSA-9006934 | 1.40E-02 | P02452;P02461;P08123;P20908;P02751;P11047 |
| *Defective HEXB causes GM2G2* | R-HSA-3656248 | 1.44E-02 | P07686 |
| *Interleukin-4 and Interleukin-13 signaling* | R-HSA-6785807 | 1.44E-02 | P08123;P08253;P02751 |
| *Antigen Presentation: Folding, assembly and peptide loading of class I MHC* | R-HSA-983170 | 1.46E-02 | P30101;P27797 |
| *Cell surface interactions at the vascular wall* | R-HSA-202733 | 1.74E-02 | P02452;P14174;P08123;P02751 |
| *GPVI-mediated activation cascade* | R-HSA-114604 | 1.85E-02 | P02452;P08123 |
| ***Keratan sulfate/keratin metabolism*** | R-HSA-1638074 | 1.91E-02 | P07686;P51884 |
| ***Glycosaminoglycan metabolism*** | R-HSA-1630316 | 1.91E-02 | P13611;P07686;P51884 |
| *Platelet degranulation* | R-HSA-114608 | 1.91E-02 | Q08380;P02751;P10909 |
| *Response to elevated platelet cytosolic Ca^2+^* | R-HSA-76005 | 1.91E-02 | Q08380;P02751;P10909 |
| *NCAM1 interactions* | R-HSA-419037 | 2.10E-02 | P02461;P20908 |
| *Diseases of glycosylation* | R-HSA-3781865 | 2.47E-02 | P13611;P07686;P51884 |
| *Immune System* | R-HSA-168256 | 2.50E-02 | P02452;P02461;P36222;P14174;P08123;P08253;Q8IV08;P07686;P30101;P02751;P27797;P10909 |
| *Chondroitin sulfate/dermatan sulfate metabolism* | R-HSA-1793185 | 2.95E-02 | P13611;P07686 |
| *Signaling by PDGF* | R-HSA-186797 | 4.17E-02 | P02461;P20908 |
| *NCAM signaling for neurite out-growth* | R-HSA-375165 | 4.72E-02 | P02461;P20908 |

| Supplementary Table 5 – Significant Reactome pathways of commonly upregulated proteins on the secretome and proteome of MSCs cultured under physioxia (long exposure) | | | |
| --- | --- | --- | --- |
| **Reactome Pathway** | **Identifier** | **FDR** | **ID mapped** |
| ***Hemostasis*** | R-HSA-109582 | 1.32E-04 | Q86UX7;P00734;P01834;P01009;P02768;P01008;P07355;Q01518 |
| *Platelet activation, signaling and aggregation* | R-HSA-76002 | 5.17E-04 | Q86UX7;P00734;P01009;P02768;Q01518 |
| *Regulation of Insulin-like Growth Factor (IGF) transport and uptake by Insulin-like Growth Factor Binding Proteins (IGFBPs)* | R-HSA-381426 | 5.17E-04 | P00734;P01009;P02768;P01008 |
| *Platelet degranulation* | R-HSA-114608 | 5.17E-04 | Q86UX7;P01009;P02768;Q01518 |
| *Response to elevated platelet cytosolic Ca^2+^* | R-HSA-76005 | 5.17E-04 | Q86UX7;P01009;P02768;Q01518 |
| *Smooth Muscle Contraction* | R-HSA-445355 | 5.17E-04 | P04083;P62736;P07355 |
| ***Scavenging of heme from plasma*** | R-HSA-2168880 | 4.77E-03 | P00738;P01834;P02768 |
| ***Post-translational protein phosphorylation*** | R-HSA-8957275 | 4.77E-03 | P01009;P02768;P01008 |
| ***Innate Immune System*** | R-HSA-168249 | 4.77E-03 | P04004;P00738;P00734;P01834;P01009;P07355;Q01518 |
| ***Common Pathway of Fibrin Clot Formation*** | R-HSA-140875 | 4.77E-03 | P00734;P01008 |
| *Intrinsic Pathway of Fibrin Clot Formation* | R-HSA-140837 | 4.80E-03 | P00734;P01008 |
| *Binding and Uptake of Ligands by Scavenger Receptors* | R-HSA-2173782 | 6.11E-03 | P00738;P01834;P02768 |
| *Regulation of Complement cascade* | R-HSA-977606 | 6.33E-03 | P04004;P00734;P01834 |
| ***Complement cascade*** | R-HSA-166658 | 7.15E-03 | P04004;P00734;P01834 |
| *Formation of Fibrin Clot (Clotting Cascade)* | R-HSA-140877 | 1.02E-02 | P00734;P01008 |
| *Muscle contraction* | R-HSA-397014 | 1.66E-02 | P04083;P62736;P07355 |
| *Neutrophil degranulation* | R-HSA-6798695 | 2.13E-02 | P00738;P01009;P07355;Q01518 |
| ***Immune System*** | R-HSA-168256 | 2.13E-02 | P04004;P00738;P00734;P04083;P01834;P01009;P07355;Q01518 |
| *Defective F8 cleavage by thrombin* | R-HSA-9672391 | 2.31E-02 | P00734 |
| *Defective factor XII causes hereditary angioedema* | R-HSA-9657688 | 2.31E-02 | P00734 |
| *Localization of the PINCH-ILK-PARVIN complex to focal adhesions* | R-HSA-446343 | 3.08E-02 | Q13418 |
| *Ciprofloxacin ADME* | R-HSA-9793528 | 3.84E-02 | P02768 |
| *Defective factor VIII causes hemophilia A* | R-HSA-9662001 | 4.48E-02 | P00734 |

| Supplementary Table 6 – Significant Reactome pathways of commonly downregulated proteins by low stiffness and oxygen levels on MSCs secretome (long exposure) | | | |
| --- | --- | --- | --- |
| **Reactome Pathway** | **Identifier** | **FDR** | **ID mapped** |
| *Assembly of collagen fibrils and other multimeric structures* | R-HSA-2022090 | 4.89E-12 | P02462;P02461;Q92626;P05997;P08123;P20908;Q99715;Q15113;P12111 |
| ***Extracellular matrix organization*** | R-HSA-1474244 | 9.10E-12 | P01033;P02462;P02461;P05997;Q92626;P35555;P20908;P12111;P11047;P21810;P08123;Q99715;Q15113 |
| ***Collagen formation*** | R-HSA-1474290 | 4.58E-11 | P02462;P02461;Q92626;P05997;P08123;P20908;Q99715;Q15113;P12111 |
| *Degradation of the extracellular matrix* | R-HSA-1474228 | 4.58E-11 | P01033;P02462;P02461;P05997;P08123;P35555;P20908;Q99715;P12111;P11047 |
| ***Collagen biosynthesis and modifying enzymes*** | R-HSA-1650814 | 1.67E-10 | P02462;P02461;P05997;P08123;P20908;Q99715;Q15113;P12111 |
| ***ECM proteoglycans*** | R-HSA-3000178 | 3.79E-10 | P02462;P02461;P05997;P21810;P08123;P20908;P12111;P11047 |
| *Collagen chain trimerization* | R-HSA-8948216 | 4.07E-10 | P02462;P02461;P05997;P08123;P20908;Q99715;P12111 |
| *Collagen degradation* | R-HSA-1442490 | 4.79E-09 | P02462;P02461;P05997;P08123;P20908;Q99715;P12111 |
| *Integrin cell surface interactions* | R-HSA-216083 | 2.93E-08 | P02462;P02461;P05997;P08123;P35555;P20908;P12111 |
| *Non-integrin membrane-ECM interactions* | R-HSA-3000171 | 1.26E-07 | P02462;P02461;P05997;P08123;P20908;P11047 |
| ***Signaling by PDGF*** | R-HSA-186797 | 1.26E-07 | P02462;P02461;P05997;P20908;P35442;P12111 |
| *MET activates PTK2 signaling* | R-HSA-8874081 | 1.89E-07 | P02461;P05997;P08123;P20908;P11047 |
| *MET promotes cell motility* | R-HSA-8875878 | 8.26E-07 | P02461;P05997;P08123;P20908;P11047 |
| *NCAM1 interactions* | R-HSA-419037 | 8.64E-07 | P02462;P02461;P05997;P20908;P12111 |
| *Crosslinking of collagen fibrils* | R-HSA-2243919 | 1.56E-06 | P02462;Q92626;P08123;Q15113 |
| *NCAM signaling for neurite out-growth* | R-HSA-375165 | 5.78E-06 | P02462;P02461;P05997;P20908;P12111 |
| *Syndecan interactions* | R-HSA-3000170 | 6.48E-06 | P02461;P05997;P08123;P20908 |
| ***Signaling by MET*** | R-HSA-6806834 | 1.56E-05 | P02461;P05997;P08123;P20908;P11047 |
| *Scavenging by Class A Receptors* | R-HSA-3000480 | 1.37E-04 | P02462;P02461;P08123 |
| ***Signaling by Receptor Tyrosine Kinases*** | R-HSA-9006934 | 3.59E-04 | P02462;P02461;P05997;P08123;P20908;P35442;P12111;P11047 |
| *Axon guidance* | R-HSA-422475 | 3.77E-04 | P02462;P02461;P05997;O00560;P20908;P12111;P11047;P62249 |
| *Nervous system development* | R-HSA-9675108 | 5.20E-04 | P02462;P02461;P05997;O00560;P20908;P12111;P11047;P62249 |
| *Post-translational protein phosphorylation* | R-HSA-8957275 | 1.11E-03 | P01033;P35555;P11047;P01034 |
| *Defective CHSY1 causes TPBS* | R-HSA-3595177 | 1.32E-03 | P21810;Q6UVK1 |
| *Defective CHST14 causes EDS, musculocontractural type* | R-HSA-3595174 | 1.32E-03 | P21810;Q6UVK1 |
| *Defective CHST3 causes SEDCJD* | R-HSA-3595172 | 1.32E-03 | P21810;Q6UVK1 |
| *Regulation of Insulin-like Growth Factor (IGF) transport and uptake by Insulin-like Growth Factor Binding Proteins (IGFBPs)* | R-HSA-381426 | 1.46E-03 | P01033;P35555;P11047;P01034 |
| *Dermatan sulfate biosynthesis* | R-HSA-2022923 | 2.12E-03 | P21810;Q6UVK1 |
| *CS/DS degradation* | R-HSA-2024101 | 3.43E-03 | P21810;Q6UVK1 |
| *Anchoring fibril formation* | R-HSA-2214320 | 3.93E-03 | P02462;P08123 |
| *Chondroitin sulfate biosynthesis* | R-HSA-2022870 | 5.77E-03 | P21810;Q6UVK1 |
| *Defective B3GALT6 causes EDSP2 and SEMDJL1* | R-HSA-4420332 | 5.77E-03 | P21810;Q6UVK1 |
| *Defective B4GALT7 causes EDS, progeroid type* | R-HSA-3560783 | 5.77E-03 | P21810;Q6UVK1 |
| *Defective B3GAT3 causes JDSSDHD* | R-HSA-3560801 | 5.77E-03 | P21810;Q6UVK1 |
| *A tetrasaccharide linker sequence is required for GAG synthesis* | R-HSA-1971475 | 9.67E-03 | P21810;Q6UVK1 |
| *Laminin interactions* | R-HSA-3000157 | 1.28E-02 | P02462;P11047 |
| *Signaling by TGFB family members* | R-HSA-9006936 | 1.42E-02 | P35555;P08123;P08476 |
| *Platelet degranulation* | R-HSA-114608 | 1.55E-02 | P01033;P07602;Q08380 |
| *Binding and Uptake of Ligands by Scavenger Receptors* | R-HSA-2173782 | 1.59E-02 | P02462;P02461;P08123 |
| *Platelet activation, signaling and aggregation* | R-HSA-76002 | 1.60E-02 | P01033;P07602;Q08380;P08123 |
| *Detoxification of Reactive Oxygen Species* | R-HSA-3299685 | 1.71E-02 | P30044-1;P30044-2 |
| *Response to elevated platelet cytosolic Ca2+* | R-HSA-76005 | 1.73E-02 | P01033;P07602;Q08380 |
| *Diseases associated with glycosaminoglycan metabolism* | R-HSA-3560782 | 1.88E-02 | P21810;Q6UVK1 |
| *Diseases of glycosylation* | R-HSA-3781865 | 2.24E-02 | P21810;Q6UVK1;P35442 |
| *Chondroitin sulfate/dermatan sulfate metabolism* | R-HSA-1793185 | 2.43E-02 | P21810;Q6UVK1 |
| *Developmental Biology* | R-HSA-1266738 | 2.43E-02 | P02462;P02461;P05997;O00560;P20908;P12111;P11047;P62249 |
| *Heparan sulfate/heparin (HS-GAG) metabolism* | R-HSA-1638091 | 2.66E-02 | P21810;Q6UVK1 |
| *Antagonism of Activin by Follistatin* | R-HSA-2473224 | 2.97E-02 | P08476 |
| ***Signal Transduction*** | R-HSA-162582 | 4.94E-02 | P07602;P02462;P02461;P05997;P08123;P35555;P20908;P08476;P35442;P02786;P12111;P11047 |

| Supplementary Table 7 – Significant Reactome pathways of inversely regulated proteins by low stiffness (up) and oxygen levels (down) on MSCs secretome (long exposure) | | | |
| --- | --- | --- | --- |
| **Reactome Pathway** | **Identifier** | **FDR** | **ID mapped** |
| ***Calnexin/calreticulin cycle*** | R-HSA-901042 | 8.51E-05 | Q14697;P30101;P27797 |
| *N-glycan trimming in the ER and Calnexin/Calreticulin cycle* | R-HSA-532668 | 1.02E-04 | Q14697;P30101;P27797 |
| *Antigen Presentation: Folding, assembly and peptide loading of class I MHC* | R-HSA-983170 | 6.36E-03 | P30101;P27797 |
| *Gluconeogenesis* | R-HSA-70263 | 6.36E-03 | P00505;P40926 |
| ***Asparagine N-linked glycosylation*** | R-HSA-446203 | 2.42E-02 | Q14697;P30101;P27797 |
| *ER-Phagosome pathway* | R-HSA-1236974 | 2.73E-02 | P30101;P27797 |
| ***Glucose metabolism*** | R-HSA-70326 | 2.84E-02 | P00505;P40926 |
| ***Antigen processing-Cross presentation*** | R-HSA-1236975 | 2.98E-02 | P30101;P27797 |
| *Maturation of spike protein* | R-HSA-9683686 | 2.99E-02 | Q14697 |
| *Scavenging by Class F Receptors* | R-HSA-3000484 | 2.99E-02 | P27797 |
| *Assembly of Viral Components at the Budding Site* | R-HSA-168316 | 2.99E-02 | P27797 |
| *ATF6 (ATF6-alpha) activates chaperone genes* | R-HSA-381183 | 3.83E-02 | P27797 |
| *Virus Assembly and Release* | R-HSA-168268 | 3.83E-02 | P27797 |
| *ATF6 (ATF6-alpha) activates chaperones* | R-HSA-381033 | 3.83E-02 | P27797 |
| *Aspartate and asparagine metabolism* | R-HSA-8963693 | 3.83E-02 | P00505 |
| *Glutamate and glutamine metabolism* | R-HSA-8964539 | 3.83E-02 | P00505 |
| *Degradation of cysteine and homocysteine* | R-HSA-1614558 | 3.83E-02 | P00505 |
| *Scavenging by Class A Receptors* | R-HSA-3000480 | 4.12E-02 | P27797 |
| *Citric acid cycle (TCA cycle)* | R-HSA-71403 | 4.12E-02 | P40926 |
| *Sulfur amino acid metabolism* | R-HSA-1614635 | 4.12E-02 | P00505 |
| *Glyoxylate metabolism and glycine degradation* | R-HSA-389661 | 4.12E-02 | P00505 |
| *Metabolism of carbohydrates* | R-HSA-71387 | 4.12E-02 | P00505;P40926 |
| *Translation of Structural Proteins* | R-HSA-9683701 | 4.12E-02 | Q14697 |
| *Signaling by high-kinase activity BRAF mutants* | R-HSA-6802948 | 4.12E-02 | P30086 |
| *Maturation of spike protein* | R-HSA-9694548 | 4.12E-02 | Q14697 |
| *Detoxification of Reactive Oxygen Species* | R-HSA-3299685 | 4.12E-02 | P30048 |
| *MAP2K and MAPK activation* | R-HSA-5674135 | 4.12E-02 | P30086 |
| *Negative regulation of MAPK pathway* | R-HSA-5675221 | 4.12E-02 | P30086 |
| *Signaling downstream of RAS mutants* | R-HSA-9649948 | 4.12E-02 | P30086 |
| *Signaling by moderate kinase activity BRAF mutants* | R-HSA-6802946 | 4.12E-02 | P30086 |
| *Paradoxical activation of RAF signaling by kinase inactive BRAF* | R-HSA-6802955 | 4.12E-02 | P30086 |
| *Signaling by RAS mutants* | R-HSA-6802949 | 4.12E-02 | P30086 |
| *Class I MHC mediated antigen processing & presentation* | R-HSA-983169 | 4.12E-02 | P30101;P27797 |
| *Pyruvate metabolism and Citric Acid (TCA) cycle* | R-HSA-71406 | 4.61E-02 | P40926 |

| *Supplementary Table 8 – Protein names of TOP 50 VIP>1 proteins represented on Figure 4A* | |
| --- | --- |
| **Accession number** | **Protein Name** |
| P98160 | Basement membrane-specific heparan sulfate proteoglycan core protein |
| P14543 | Nidogen-1 |
| Q14112 | Nidogen-2 |
| P20908 | Collagen alpha-1(V) chain |
| Q9Y6C2 | EMILIN-1 |
| P08572 | Collagen alpha-2(IV) chain |
| P09871 | Complement C1s subcomponent |
| P04004 | Vitronectin |
| P02452 | Collagen alpha-1(I) chain |
| P08476 | Inhibin beta A chain |
| Q15063 | Periostin |
| P00441 | Superoxide dismutase [Cu-Zn] |
| P35556 | Fibrillin-2 |
| P07585 | Decorin |
| P21810 | Biglycan |
| P28799 | Progranulin |
| P08123 | Collagen alpha-2(I) chain |
| P19022 | Cadherin-2 |
| P02743 | Serum amyloid P-component |
| P12110 | Collagen alpha-2(VI) chain |
| P12109 | Collagen alpha-1(VI) chain |
| P05546 | Heparin cofactor 2 |
| Q92859 | Neogenin |
| P16035 | Metalloproteinase inhibitor 2 |
| Q9NRN5 | Olfactomedin-like protein 3 |
| P02747 | Complement C1q subcomponent subunit C |
| P10909 | Clusterin |
| Q8NBJ4 | Golgi membrane protein 1 |
| Q92626 | Peroxidasin homolog |
| P00736 | Complement C1r subcomponent |
| O95445 | Apolipoprotein M |
| P12111 | Collagen alpha-3(VI) chain |
| P61916 | NPC intracellular cholesterol transporter 2 |
| P07858 | Cathepsin B |
| Q9Y3Q0 | N-acetylated-alpha-linked acidic dipeptidase 2 |
| P02461 | Collagen alpha-1(III) chain |
| P55268 | Laminin subunit beta-2 |
| Q96CG8 | Collagen triple helix repeat-containing protein 1 |
| Q9Y240 | C-type lectin domain family 11 member A |
| Q92743 | Serine protease HTRA1 |
| P01034 | Cystatin-C |
| P18206 | Vinculin |
| O60462 | Neuropilin-2 |
| P07339 | Cathepsin D |
| P02751 | Fibronectin |
| P09382 | Galectin-1 |
| P01859 | Immunoglobulin heavy constant gamma 2 |
| P02790 | Hemopexin |
| P07711 | Procathepsin L |
| P01024 | Complement C3 |

| Supplementary Table 9 – Significant Reactome pathways of downregulated proteins by physiological priming on MSCs secretome | | | |
| --- | --- | --- | --- |
| **Reactome Pathway** | **Identifier** | **FDR** | **ID mapped** |
| **Extracellular matrix organization** | R-HSA-1474244 | 6.03E-09 | P02462;P23142;P03956;Q14767;P35555;P12107;Q02809;P13611;P05121;Q99715;P51884 |
| Regulation of Insulin-like Growth Factor (IGF) transport and uptake by Insulin-like Growth Factor Binding Proteins (IGFBPs) | R-HSA-381426 | 7.41E-04 | O43852;Q13219;P03956;P35555;P13611 |
| Collagen degradation | R-HSA-1442490 | 7.41E-04 | P02462;P03956;P12107;Q99715 |
| Degradation of the extracellular matrix | R-HSA-1474228 | 7.41E-04 | P02462;P03956;P35555;P12107;Q99715 |
| Collagen biosynthesis and modifying enzymes | R-HSA-1650814 | 7.41E-04 | P02462;Q02809;P12107;Q99715 |
| **ECM proteoglycans** | R-HSA-3000178 | 9.89E-04 | P02462;P13611;P05121;P51884 |
| Collagen formation | R-HSA-1474290 | 1.64E-03 | P02462;Q02809;P12107;Q99715 |
| Molecules associated with elastic fibres | R-HSA-2129379 | 2.34E-03 | P23142;Q14767;P35555 |
| Diseases associated with glycosaminoglycan metabolism | R-HSA-3560782 | 2.82E-03 | P13611;Q16394;P51884 |
| Elastic fibre formation | R-HSA-1566948 | 2.89E-03 | P23142;Q14767;P35555 |
| Collagen chain trimerization | R-HSA-8948216 | 2.89E-03 | P02462;P12107;Q99715 |
| Platelet degranulation | R-HSA-114608 | 3.67E-03 | P11021;O43852;Q08380;P05121 |
| Response to elevated platelet cytosolic Ca2+ | R-HSA-76005 | 3.77E-03 | P11021;O43852;Q08380;P05121 |
| HDL remodeling | R-HSA-8964058 | 4.56E-03 | P55058-1;P55058-2 |
| Diseases of glycosylation | R-HSA-3781865 | 4.69E-03 | P13611;P35442;Q16394;P51884 |
| Assembly of collagen fibrils and other multimeric structures | R-HSA-2022090 | 4.87E-03 | P02462;P12107;Q99715 |
| Integrin cell surface interactions | R-HSA-216083 | 1.17E-02 | P02462;P35555;P51884 |
| Signaling by TGF-beta Receptor Complex | R-HSA-170834 | 1.57E-02 | Q14767;P35555;P05121 |
| Post-translational protein phosphorylation | R-HSA-8957275 | 2.07E-02 | O43852;P35555;P13611 |
| Diseases of metabolism | R-HSA-5668914 | 2.69E-02 | P13611;P35442;Q16394;P51884 |
| Signaling by TGFB family members | R-HSA-9006936 | 2.69E-02 | Q14767;P35555;P05121 |
| Plasma lipoprotein remodeling | R-HSA-8963899 | 2.69E-02 | P55058-1;P55058-2 |
| Glycosaminoglycan metabolism | R-HSA-1630316 | 2.69E-02 | P13611;Q16394;P51884 |
| Platelet activation, signaling and aggregation | R-HSA-76002 | 2.72E-02 | P11021;O43852;Q08380;P05121 |
| NR1H3 & NR1H2 regulate gene expression linked to cholesterol transport and efflux | R-HSA-9029569 | 2.79E-02 | P55058-1;P55058-2 |
| TGF-beta receptor signaling activates SMADs | R-HSA-2173789 | 3.85E-02 | Q14767;P35555 |
| Hemostasis | R-HSA-109582 | 3.85E-02 | P11021;O43852;Q08380;P03956;Q71U36;P05121 |
| NR1H2 and NR1H3-mediated signaling | R-HSA-9024446 | 3.85E-02 | P55058-1;P55058-2 |

| Supplementary Table 10 – Significant Reactome pathways of common downregulated synthetized and secreted proteins by MSCs upon mechanomodulation priming | | | |
| --- | --- | --- | --- |
| **Reactome Pathway** | **Identifier** | **FDR** | **ID mapped** |
| ECM proteoglycans | R-HSA-3000178 | 2.09E-14 | P02461;P12110;P55268;P12109;P08123;P20908;P13611;P12111;P51884 |
| **Extracellular matrix organization** | R-HSA-1474244 | 2.09E-14 | Q9Y6C2;P23142;P12110;P02461;P55268;P12109;P08123;P20908;P13611;Q99715;P12111;P51884 |
| Collagen chain trimerization | R-HSA-8948216 | 6.11E-12 | P02461;P12110;P12109;P08123;P20908;Q99715;P12111 |
| Assembly of collagen fibrils and other multimeric structures | R-HSA-2022090 | 4.40E-11 | P02461;P12110;P12109;P08123;P20908;Q99715;P12111 |
| Collagen degradation | R-HSA-1442490 | 4.81E-11 | P02461;P12110;P12109;P08123;P20908;Q99715;P12111 |
| Collagen biosynthesis and modifying enzymes | R-HSA-1650814 | 5.51E-11 | P02461;P12110;P12109;P08123;P20908;Q99715;P12111 |
| Integrin cell surface interactions | R-HSA-216083 | 2.49E-10 | P02461;P12110;P12109;P08123;P20908;P12111;P51884 |
| Collagen formation | R-HSA-1474290 | 3.14E-10 | P02461;P12110;P12109;P08123;P20908;Q99715;P12111 |
| Degradation of the extracellular matrix | R-HSA-1474228 | 6.05E-09 | P02461;P12110;P12109;P08123;P20908;Q99715;P12111 |
| NCAM1 interactions | R-HSA-419037 | 3.17E-08 | P02461;P12110;P12109;P20908;P12111 |
| **Signaling by PDGF** | R-HSA-186797 | 1.65E-07 | P02461;P12110;P12109;P20908;P12111 |
| NCAM signaling for neurite out-growth | R-HSA-375165 | 1.99E-07 | P02461;P12110;P12109;P20908;P12111 |
| MET activates PTK2 signaling | R-HSA-8874081 | 6.95E-07 | P02461;P55268;P08123;P20908 |
| MET promotes cell motility | R-HSA-8875878 | 2.06E-06 | P02461;P55268;P08123;P20908 |
| **Signaling by Receptor Tyrosine Kinases** | R-HSA-9006934 | 2.36E-06 | P02461;P12110;P55268;P12109;P08123;P20908;O60462;P12111 |
| Non-integrin membrane-ECM interactions | R-HSA-3000171 | 7.24E-06 | P02461;P55268;P08123;P20908 |
| **Signaling by MET** | R-HSA-6806834 | 2.40E-05 | P02461;P55268;P08123;P20908 |
| Syndecan interactions | R-HSA-3000170 | 4.05E-05 | P02461;P08123;P20908 |
| Diseases associated with glycosaminoglycan metabolism | R-HSA-3560782 | 1.12E-04 | Q6UVK1;P13611;P51884 |
| Defective CHSY1 causes TPBS | R-HSA-3595177 | 2.51E-04 | Q6UVK1;P13611 |
| Defective CHST3 causes SEDCJD | R-HSA-3595172 | 2.51E-04 | Q6UVK1;P13611 |
| Defective CHST14 causes EDS, musculocontractural type | R-HSA-3595174 | 2.51E-04 | Q6UVK1;P13611 |
| Axon guidance | R-HSA-422475 | 3.51E-04 | P02461;P12110;P12109;P20908;O60462;P12111 |
| Nervous system development | R-HSA-9675108 | 3.51E-04 | P02461;P12110;P12109;P20908;O60462;P12111 |
| Dermatan sulfate biosynthesis | R-HSA-2022923 | 3.56E-04 | Q6UVK1;P13611 |
| CS/DS degradation | R-HSA-2024101 | 5.75E-04 | Q6UVK1;P13611 |
| Scavenging by Class A Receptors | R-HSA-3000480 | 1.05E-03 | P02461;P08123 |
| Chondroitin sulfate biosynthesis | R-HSA-2022870 | 1.17E-03 | Q6UVK1;P13611 |
| Defective B4GALT7 causes EDS, progeroid type | R-HSA-3560783 | 1.17E-03 | Q6UVK1;P13611 |
| Defective B3GALT6 causes EDSP2 and SEMDJL1 | R-HSA-4420332 | 1.17E-03 | Q6UVK1;P13611 |
| Defective B3GAT3 causes JDSSDHD | R-HSA-3560801 | 1.17E-03 | Q6UVK1;P13611 |
| A tetrasaccharide linker sequence is required for GAG synthesis | R-HSA-1971475 | 1.31E-03 | Q6UVK1;P13611 |
| Glycosaminoglycan metabolism | R-HSA-1630316 | 1.51E-03 | Q6UVK1;P13611;P51884 |
| Diseases of glycosylation | R-HSA-3781865 | 2.30E-03 | Q6UVK1;P13611;P51884 |
| Molecules associated with elastic fibres | R-HSA-2129379 | 2.62E-03 | Q9Y6C2;P23142 |
| Elastic fibre formation | R-HSA-1566948 | 3.69E-03 | Q9Y6C2;P23142 |
| Chondroitin sulfate/dermatan sulfate metabolism | R-HSA-1793185 | 4.74E-03 | Q6UVK1;P13611 |
| Heparan sulfate/heparin (HS-GAG) metabolism | R-HSA-1638091 | 6.12E-03 | Q6UVK1;P13611 |
| Defective VWF binding to collagen type I | R-HSA-9845622 | 8.68E-03 | P08123 |
| Diseases of metabolism | R-HSA-5668914 | 1.01E-02 | Q6UVK1;P13611;P51884 |
| Defective VWF cleavage by ADAMTS13 variant | R-HSA-9845621 | 1.01E-02 | P08123 |
| Enhanced cleavage of VWF variant by ADAMTS13 | R-HSA-9845619 | 1.01E-02 | P08123 |
| Neurophilin interactions with VEGF and VEGFR | R-HSA-194306 | 1.01E-02 | O60462 |
| Amyloid fiber formation | R-HSA-977225 | 1.01E-02 | Q15582;P02743 |
| Developmental Biology | R-HSA-1266738 | 1.01E-02 | P02461;P12110;P12109;P20908;O60462;P12111 |
| **Metabolism of carbohydrates** | R-HSA-71387 | 1.01E-02 | Q6UVK1;P13611;P51884 |
| Enhanced binding of GP1BA variant to VWF multimer:collagen | R-HSA-9845620 | 1.01E-02 | P08123 |
| Defective binding of VWF variant to GPIb:IX:V | R-HSA-9846298 | 1.01E-02 | P08123 |
| NrCAM interactions | R-HSA-447038 | 1.01E-02 | O60462 |
| Post-translational protein phosphorylation | R-HSA-8957275 | 1.03E-02 | P55268;P13611 |
| Defects of platelet adhesion to exposed collagen | R-HSA-9823587 | 1.15E-02 | P08123 |
| Defective B4GALT1 causes B4GALT1-CDG (CDG-2d) | R-HSA-3656244 | 1.15E-02 | P51884 |
| Defective ST3GAL3 causes MCT12 and EIEE15 | R-HSA-3656243 | 1.15E-02 | P51884 |
| Defective CHST6 causes MCDC1 | R-HSA-3656225 | 1.15E-02 | P51884 |
| Regulation of Insulin-like Growth Factor (IGF) transport and uptake by Insulin-like Growth Factor Binding Proteins (IGFBPs) | R-HSA-381426 | 1.37E-02 | P55268;P13611 |
| Binding and Uptake of Ligands by Scavenger Receptors | R-HSA-2173782 | 1.47E-02 | P02461;P08123 |
| GP1b-IX-V activation signalling | R-HSA-430116 | 1.73E-02 | P08123 |
| **Signal Transduction** | R-HSA-162582 | 2.06E-02 | P02461;P12110;P55268;P12109;P08123;P20908;O60462;P12111 |
| Keratan sulfate degradation | R-HSA-2022857 | 2.15E-02 | P51884 |
| Anchoring fibril formation | R-HSA-2214320 | 2.15E-02 | P08123 |
| Platelet Adhesion to exposed collagen | R-HSA-75892 | 2.29E-02 | P08123 |
| Crosslinking of collagen fibrils | R-HSA-2243919 | 2.58E-02 | P08123 |
| Diseases of hemostasis | R-HSA-9671793 | 2.72E-02 | P08123 |
| Keratan sulfate biosynthesis | R-HSA-2022854 | 3.98E-02 | P51884 |
| Laminin interactions | R-HSA-3000157 | 4.26E-02 | P55268 |
| Immunoregulatory interactions between a Lymphoid and a non-Lymphoid cell | R-HSA-198933 | 4.30E-02 | P02461;P08123 |
| GPVI-mediated activation cascade | R-HSA-114604 | 4.95E-02 | P08123 |

| Supplementary Table 11 – Significant Reactome pathways of common downregulated secreted proteins by mechanomodulated MSCs when cultured on soft platforms for long (7-10 days) or short (48h) periods | | | |
| --- | --- | --- | --- |
| **Reactome Pathway** | **Identifier** | **FDR** | **ID mapped** |
| **Extracellular matrix organization** | R-HSA-1474244 | 4.06E-14 | P02462;P02461;P05997;Q92626;Q14767;P35555;P20908;P12111;P11047;P12110;P21810;P08123;P05121;Q99715 |
| ECM proteoglycans | R-HSA-3000178 | 4.06E-14 | P02462;P02461;P12110;P05997;P21810;P08123;P20908;P05121;P12111;P11047 |
| Assembly of collagen fibrils and other multimeric structures | R-HSA-2022090 | 3.20E-13 | P02462;P02461;P12110;Q92626;P05997;P08123;P20908;Q99715;P12111 |
| Collagen chain trimerization | R-HSA-8948216 | 1.73E-12 | P02462;P02461;P12110;P05997;P08123;P20908;Q99715;P12111 |
| Collagen formation | R-HSA-1474290 | 5.25E-12 | P02462;P02461;P12110;Q92626;P05997;P08123;P20908;Q99715;P12111 |
| Degradation of the extracellular matrix | R-HSA-1474228 | 5.25E-12 | P02462;P02461;P12110;P05997;P08123;P35555;P20908;Q99715;P12111;P11047 |
| Collagen degradation | R-HSA-1442490 | 1.91E-11 | P02462;P02461;P12110;P05997;P08123;P20908;Q99715;P12111 |
| Collagen biosynthesis and modifying enzymes | R-HSA-1650814 | 2.43E-11 | P02462;P02461;P12110;P05997;P08123;P20908;Q99715;P12111 |
| Integrin cell surface interactions | R-HSA-216083 | 1.38E-10 | P02462;P02461;P12110;P05997;P08123;P35555;P20908;P12111 |
| **Signaling by PDGF** | R-HSA-186797 | 6.65E-10 | P02462;P02461;P12110;P05997;P20908;P35442;P12111 |
| NCAM1 interactions | R-HSA-419037 | 5.03E-09 | P02462;P02461;P12110;P05997;P20908;P12111 |
| Non-integrin membrane-ECM interactions | R-HSA-3000171 | 3.42E-08 | P02462;P02461;P05997;P08123;P20908;P11047 |
| NCAM signaling for neurite out-growth | R-HSA-375165 | 4.99E-08 | P02462;P02461;P12110;P05997;P20908;P12111 |
| MET activates PTK2 signaling | R-HSA-8874081 | 5.51E-08 | P02461;P05997;P08123;P20908;P11047 |
| MET promotes cell motility | R-HSA-8875878 | 2.59E-07 | P02461;P05997;P08123;P20908;P11047 |
| Syndecan interactions | R-HSA-3000170 | 2.82E-06 | P02461;P05997;P08123;P20908 |
| Signaling by MET | R-HSA-6806834 | 6.04E-06 | P02461;P05997;P08123;P20908;P11047 |
| Signaling by Receptor Tyrosine Kinases | R-HSA-9006934 | 1.01E-05 | P02462;P02461;P12110;P05997;P08123;P20908;P35442;P12111;P11047 |
| Signaling by TGFB family members | R-HSA-9006936 | 4.33E-05 | Q14767;P35555;P08123;P08476;P05121 |
| Crosslinking of collagen fibrils | R-HSA-2243919 | 5.38E-05 | P02462;Q92626;P08123 |
| Scavenging by Class A Receptors | R-HSA-3000480 | 5.38E-05 | P02462;P02461;P08123 |
| Signaling by TGF-beta Receptor Complex | R-HSA-170834 | 2.68E-04 | Q14767;P35555;P08123;P05121 |
| Diseases associated with glycosaminoglycan metabolism | R-HSA-3560782 | 5.23E-04 | P21810;Q6UVK1;Q16394 |
| Defective CHSY1 causes TPBS | R-HSA-3595177 | 5.98E-04 | P21810;Q6UVK1 |
| Defective CHST14 causes EDS, musculocontractural type | R-HSA-3595174 | 5.98E-04 | P21810;Q6UVK1 |
| Defective CHST3 causes SEDCJD | R-HSA-3595172 | 5.98E-04 | P21810;Q6UVK1 |
| Regulation of Insulin-like Growth Factor (IGF) transport and uptake by Insulin-like Growth Factor Binding Proteins (IGFBPs) | R-HSA-381426 | 6.21E-04 | P24592;P35555;P11047;P01034 |
| Axon guidance | R-HSA-422475 | 6.51E-04 | P02462;P02461;P12110;P05997;P20908;P12111;P11047 |
| Nervous system development | R-HSA-9675108 | 8.45E-04 | P02462;P02461;P12110;P05997;P20908;P12111;P11047 |
| Heparan sulfate/heparin (HS-GAG) metabolism | R-HSA-1638091 | 8.45E-04 | P21810;Q6UVK1;Q16394 |
| Dermatan sulfate biosynthesis | R-HSA-2022923 | 8.45E-04 | P21810;Q6UVK1 |
| Diseases of glycosylation | R-HSA-3781865 | 8.65E-04 | P21810;Q6UVK1;P35442;Q16394 |
| **Signal Transduction** | R-HSA-162582 | 1.30E-03 | P02462;P02461;P05997;Q8IVT5;Q14767;P35555;P20908;P35442;P12111;P11047;P12110;P08123;P08476;P05121 |
| CS/DS degradation | R-HSA-2024101 | 1.36E-03 | P21810;Q6UVK1 |
| Anchoring fibril formation | R-HSA-2214320 | 1.56E-03 | P02462;P08123 |
| Chondroitin sulfate biosynthesis | R-HSA-2022870 | 2.76E-03 | P21810;Q6UVK1 |
| Defective B4GALT7 causes EDS, progeroid type | R-HSA-3560783 | 2.76E-03 | P21810;Q6UVK1 |
| Defective B3GALT6 causes EDSP2 and SEMDJL1 | R-HSA-4420332 | 2.76E-03 | P21810;Q6UVK1 |
| Defective B3GAT3 causes JDSSDHD | R-HSA-3560801 | 2.76E-03 | P21810;Q6UVK1 |
| A tetrasaccharide linker sequence is required for GAG synthesis | R-HSA-1971475 | 3.08E-03 | P21810;Q6UVK1 |
| Post-translational protein phosphorylation | R-HSA-8957275 | 3.38E-03 | P35555;P11047;P01034 |
| Laminin interactions | R-HSA-3000157 | 4.08E-03 | P02462;P11047 |
| Diseases of metabolism | R-HSA-5668914 | 4.43E-03 | P21810;Q6UVK1;P35442;Q16394 |
| Glycosaminoglycan metabolism | R-HSA-1630316 | 5.36E-03 | P21810;Q6UVK1;Q16394 |
| Binding and Uptake of Ligands by Scavenger Receptors | R-HSA-2173782 | 5.73E-03 | P02462;P02461;P08123 |
| SMAD2/SMAD3:SMAD4 heterotrimer regulates transcription | R-HSA-2173796 | 5.83E-03 | P08123;P05121 |
| Molecules associated with elastic fibres | R-HSA-2129379 | 6.15E-03 | Q14767;P35555 |
| Elastic fibre formation | R-HSA-1566948 | 8.62E-03 | Q14767;P35555 |
| TGF-beta receptor signaling activates SMADs | R-HSA-2173789 | 9.79E-03 | Q14767;P35555 |
| Chondroitin sulfate/dermatan sulfate metabolism | R-HSA-1793185 | 1.10E-02 | P21810;Q6UVK1 |
| Transcriptional activity of SMAD2/SMAD3:SMAD4 heterotrimer | R-HSA-2173793 | 1.15E-02 | P08123;P05121 |
| Defective VWF binding to collagen type I | R-HSA-9845622 | 1.33E-02 | P08123 |
| Antagonism of Activin by Follistatin | R-HSA-2473224 | 1.69E-02 | P08476 |
| Defective VWF cleavage by ADAMTS13 variant | R-HSA-9845621 | 1.69E-02 | P08123 |
| Enhanced cleavage of VWF variant by ADAMTS13 | R-HSA-9845619 | 1.69E-02 | P08123 |
| Amyloid fiber formation | R-HSA-977225 | 1.69E-02 | Q15582;P01034 |
| Defective binding of VWF variant to GPIb:IX:V | R-HSA-9846298 | 1.69E-02 | P08123 |
| Enhanced binding of GP1BA variant to VWF multimer:collagen | R-HSA-9845620 | 1.69E-02 | P08123 |
| Developmental Biology | R-HSA-1266738 | 1.69E-02 | P02462;P02461;P12110;P05997;P20908;P12111;P11047 |
| Defects of platelet adhesion to exposed collagen | R-HSA-9823587 | 1.76E-02 | P08123 |
| Platelet activation, signaling and aggregation | R-HSA-76002 | 2.04E-02 | Q08380;P08123;P05121 |
| Glycoprotein hormones | R-HSA-209822 | 2.19E-02 | P08476 |
| GP1b-IX-V activation signalling | R-HSA-430116 | 2.63E-02 | P08123 |
| Peptide hormone biosynthesis | R-HSA-209952 | 2.63E-02 | P08476 |
| Dissolution of Fibrin Clot | R-HSA-75205 | 2.84E-02 | P05121 |
| Metabolism of carbohydrates | R-HSA-71387 | 2.90E-02 | P21810;Q6UVK1;Q16394 |
| Defective EXT1 causes exostoses 1, TRPS2 and CHDS | R-HSA-3656253 | 3.06E-02 | Q16394 |
| Defective EXT2 causes exostoses 2 | R-HSA-3656237 | 3.06E-02 | Q16394 |
| Platelet degranulation | R-HSA-114608 | 3.26E-02 | Q08380;P05121 |
| Platelet Adhesion to exposed collagen | R-HSA-75892 | 3.49E-02 | P08123 |
| Response to elevated platelet cytosolic Ca2+ | R-HSA-76005 | 3.49E-02 | Q08380;P05121 |
| Signaling by Activin | R-HSA-1502540 | 3.70E-02 | P08476 |
| Diseases of hemostasis | R-HSA-9671793 | 4.13E-02 | P08123 |

| Supplementary Table 12 – Significant Reactome pathways of common upregulated secreted proteins by physioxia MSCs when cultured under physioxia environments for long (7-10 days) or short (48h) periods | | | |
| --- | --- | --- | --- |
| **Reactome Pathway** | **Identifier** | **FDR** | **ID mapped** |
| **Regulation of Insulin-like Growth Factor (IGF) transport and uptake by Insulin-like Growth Factor Binding Proteins (IGFBPs)** | R-HSA-381426 | 8.86E-08 | P00450;P05546;P01042;P00747;P02649;P02647;P01024 |
| **Post-translational protein phosphorylation** | R-HSA-8957275 | 9.33E-07 | P00450;P05546;P01042;P02649;P02647;P01024 |
| Binding and Uptake of Ligands by Scavenger Receptors | R-HSA-2173782 | 1.87E-06 | P04433;P00738;P68871;P02649;P02647;P80748 |
| Scavenging of heme from plasma | R-HSA-2168880 | 1.40E-05 | P04433;P00738;P68871;P02647;P80748 |
| Hemostasis | R-HSA-109582 | 1.40E-05 | P04433;P05546;P01042;P00747;P68871;P02647;P04196;P80748;P02763 |
| Retinoid metabolism and transport | R-HSA-975634 | 1.40E-05 | O95445;P02649;P02647;P02766 |
| Metabolism of fat-soluble vitamins | R-HSA-6806667 | 1.66E-05 | O95445;P02649;P02647;P02766 |
| Platelet degranulation | R-HSA-114608 | 2.22E-05 | P01042;P00747;P02647;P04196;P02763 |
| Response to elevated platelet cytosolic Ca^2+^ | R-HSA-76005 | 2.47E-05 | P01042;P00747;P02647;P04196;P02763 |
| Regulation of Complement cascade | R-HSA-977606 | 2.47E-05 | P04433;P02748;P02747;P80748;P01024 |
| **Complement cascade** | R-HSA-166658 | 3.31E-05 | P04433;P02748;P02747;P80748;P01024 |
| **Innate Immune System** | R-HSA-168249 | 4.74E-05 | P04433;P00738;P68871;P02748;P02747;P02766;P80748;P02763;P01024;P02750 |
| Visual phototransduction | R-HSA-2187338 | 1.66E-04 | O95445;P02649;P02647;P02766 |
| Initial triggering of complement | R-HSA-166663 | 2.22E-04 | P04433;P02747;P80748;P01024 |
| Platelet activation, signaling and aggregation | R-HSA-76002 | 4.20E-04 | P01042;P00747;P02647;P04196;P02763 |
| Neutrophil degranulation | R-HSA-6798695 | 5.50E-04 | P00738;P68871;P02766;P02763;P01024;P02750 |
| Chylomicron remodeling | R-HSA-8963901 | 8.62E-04 | P02649;P02647 |
| Chylomicron assembly | R-HSA-8963888 | 8.62E-04 | P02649;P02647 |
| HDL remodeling | R-HSA-8964058 | 8.93E-04 | P02649;P02647 |
| Dissolution of Fibrin Clot | R-HSA-75205 | 1.25E-03 | P00747;P04196 |
| Metabolism of vitamins and cofactors | R-HSA-196854 | 1.46E-03 | O95445;P02649;P02647;P02766 |
| Amyloid fiber formation | R-HSA-977225 | 1.47E-03 | P02649;P02647;P02766 |
| Plasma lipoprotein assembly | R-HSA-8963898 | 2.20E-03 | P02649;P02647 |
| Scavenging by Class A Receptors | R-HSA-3000480 | 2.20E-03 | P02649;P02647 |
| Classical antibody-mediated complement activation | R-HSA-173623 | 2.34E-03 | P04433;P02747;P80748 |
| Creation of C4 and C2 activators | R-HSA-166786 | 2.36E-03 | P04433;P02747;P80748 |
| Intrinsic Pathway of Fibrin Clot Formation | R-HSA-140837 | 2.57E-03 | P05546;P01042 |
| **Vesicle-mediated transport** | R-HSA-5653656 | 3.90E-03 | P04433;P00738;P68871;P02649;P02647;P80748 |
| **Immune System** | R-HSA-168256 | 4.03E-03 | P04433;P00738;P68871;P02748;P02747;P02766;P80748;P02763;P01024;P02750 |
| Plasma lipoprotein remodeling | R-HSA-8963899 | 5.89E-03 | P02649;P02647 |
| Plasma lipoprotein clearance | R-HSA-8964043 | 6.57E-03 | P02649;P02647 |
| Formation of Fibrin Clot (Clotting Cascade) | R-HSA-140877 | 7.29E-03 | P05546;P01042 |
| Heme signaling | R-HSA-9707616 | 7.87E-03 | P68871;P02647 |
| Metabolism of proteins | R-HSA-392499 | 7.95E-03 | P00450;P05546;P01042;P00747;P02649;P01019;P02647;P02766;P01024 |
| Defective SLC40A1 causes hemochromatosis 4 (HFE4) (macrophages) | R-HSA-5619049 | 9.71E-03 | P00450 |
| Defective ABCA1 causes TGD | R-HSA-5682113 | 9.71E-03 | P02647 |
| Defective CP causes aceruloplasminemia (ACERULOP) | R-HSA-5619060 | 9.71E-03 | P00450 |
| Peptide ligand-binding receptors | R-HSA-375276 | 1.14E-02 | P01042;P01019;P01024 |
| CD22 mediated BCR regulation | R-HSA-5690714 | 1.61E-02 | P04433;P80748 |
| Immunoregulatory interactions between a Lymphoid and a non-Lymphoid cell | R-HSA-198933 | 1.61E-02 | P04433;P80748;P01024 |
| Plasma lipoprotein assembly, remodeling, and clearance | R-HSA-174824 | 1.61E-02 | P02649;P02647 |
| Parasitic Infection Pathways | R-HSA-9824443 | 1.61E-02 | P04433;P80748;P01024 |
| Leishmania infection | R-HSA-9658195 | 1.61E-02 | P04433;P80748;P01024 |
| Alternative complement activation | R-HSA-173736 | 1.61E-02 | P01024 |
| HDL clearance | R-HSA-8964011 | 1.61E-02 | P02647 |
| Chylomicron clearance | R-HSA-8964026 | 1.61E-02 | P02649 |
| Scavenging by Class B Receptors | R-HSA-3000471 | 1.94E-02 | P02647 |
| Antigen activates B Cell Receptor (BCR) leading to generation of second messengers | R-HSA-983695 | 2.05E-02 | P04433;P80748 |
| Activation of C3 and C5 | R-HSA-174577 | 2.09E-02 | P01024 |
| FCGR activation | R-HSA-2029481 | 2.09E-02 | P04433;P80748 |
| Role of LAT2/NTAL/LAB on calcium mobilization | R-HSA-2730905 | 2.09E-02 | P04433;P80748 |
| Terminal pathway of complement | R-HSA-166665 | 2.09E-02 | P02748 |
| HDL assembly | R-HSA-8963896 | 2.09E-02 | P02647 |
| Erythrocytes take up oxygen and release carbon dioxide | R-HSA-1247673 | 2.09E-02 | P68871 |
| G alpha (i) signalling events | R-HSA-418594 | 2.09E-02 | P01042;P01019;P01024 |
| Role of phospholipids in phagocytosis | R-HSA-2029485 | 2.09E-02 | P04433;P80748 |
| FCERI mediated Ca^2+^ mobilization | R-HSA-2871809 | 2.09E-02 | P04433;P80748 |
| PPARA activates gene expression | R-HSA-1989781 | 2.09E-02 | P01019;P02647 |
| FCERI mediated MAPK activation | R-HSA-2871796 | 2.09E-02 | P04433;P80748 |
| Class A/1 (Rhodopsin-like receptors) | R-HSA-373076 | 2.09E-02 | P01042;P01019;P01024 |
| Regulation of lipid metabolism by PPARalpha | R-HSA-400206 | 2.09E-02 | P01019;P02647 |
| Sensory Perception | R-HSA-9709957 | 2.09E-02 | O95445;P02649;P02647;P02766 |
| FCGR3A-mediated IL10 synthesis | R-HSA-9664323 | 2.09E-02 | P04433;P80748 |
| Erythrocytes take up carbon dioxide and release oxygen | R-HSA-1237044 | 2.09E-02 | P68871 |
| O2/CO2 exchange in erythrocytes | R-HSA-1480926 | 2.09E-02 | P68871 |
| Retinoid cycle disease events | R-HSA-2453864 | 2.09E-02 | P02766 |
| Diseases of the neuronal system | R-HSA-9675143 | 2.09E-02 | P02766 |
| Diseases associated with visual transduction | R-HSA-2474795 | 2.09E-02 | P02766 |
| Post-translational protein modification | R-HSA-597592 | 2.31E-02 | P00450;P05546;P01042;P02649;P02647;P01024 |
| Leishmania phagocytosis | R-HSA-9664417 | 2.39E-02 | P04433;P80748 |
| FCGR3A-mediated phagocytosis | R-HSA-9664422 | 2.39E-02 | P04433;P80748 |
| Parasite infection | R-HSA-9664407 | 2.39E-02 | P04433;P80748 |
| Regulation of actin dynamics for phagocytic cup formation | R-HSA-2029482 | 2.42E-02 | P04433;P80748 |
| Transport of small molecules | R-HSA-382551 | 2.84E-02 | P00450;P02649;P68871;P02647 |
| Potential therapeutics for SARS | R-HSA-9679191 | 2.86E-02 | P04433;P80748 |
| Metabolism of Angiotensinogen to Angiotensins | R-HSA-2022377 | 2.88E-02 | P01019 |
| ABC transporters in lipid homeostasis | R-HSA-1369062 | 2.88E-02 | P02647 |
| FCERI mediated NF-kB activation | R-HSA-2871837 | 2.95E-02 | P04433;P80748 |
| Leishmania parasite growth and survival | R-HSA-9664433 | 2.99E-02 | P04433;P80748 |
| Anti-inflammatory response favouring Leishmania parasite infection | R-HSA-9662851 | 2.99E-02 | P04433;P80748 |
| Fcgamma receptor (FCGR) dependent phagocytosis | R-HSA-2029480 | 3.22E-02 | P04433;P80748 |
| Signaling by the B Cell Receptor (BCR) | R-HSA-983705 | 3.25E-02 | P04433;P80748 |
| Disorders of transmembrane transporters | R-HSA-5619115 | 3.42E-02 | P00450;P02647 |
| Chaperone Mediated Autophagy | R-HSA-9613829 | 3.50E-02 | P68871 |
| Common Pathway of Fibrin Clot Formation | R-HSA-140875 | 3.50E-02 | P05546 |
| The canonical retinoid cycle in rods (twilight vision) | R-HSA-2453902 | 3.66E-02 | P02766 |
| Metal ion SLC transporters | R-HSA-425410 | 3.82E-02 | P00450 |
| GPCR ligand binding | R-HSA-500792 | 3.86E-02 | P01042;P01019;P01024 |
| Purinergic signaling in leishmaniasis infection | R-HSA-9660826 | 4.29E-02 | P01024 |
| Cell recruitment (pro-inflammatory response) | R-HSA-9664424 | 4.29E-02 | P01024 |
| Fc epsilon receptor (FCERI) signaling | R-HSA-2454202 | 4.79E-02 | P04433;P80748 |
| G alpha (q) signalling events | R-HSA-416476 | 4.87E-02 | P01042;P01019 |

| Supplementary Table 13 – Significant Reactome pathways of common downregulated secreted proteins by physioxia MSCs when cultured under physioxia environments for long (7-10 days) or short (48h) periods | | | |
| --- | --- | --- | --- |
| **Reactome Pathway** | **Identifier** | **FDR** | **ID mapped** |
| **Extracellular matrix organization** | 3.79E-09 | 3.79E-09 | O60568;P02462;P23142;P07339;P35555;P13611;P08253;Q99715;Q15113;P51884 |
| Degradation of the extracellular matrix | 1.71E-04 | 1.71E-04 | P02462;P07339;P35555;P08253;Q99715 |
| Collagen degradation | 1.71E-04 | 1.71E-04 | P02462;P07339;P08253;Q99715 |
| Collagen biosynthesis and modifying enzymes | 1.71E-04 | 1.71E-04 | O60568;P02462;Q99715;Q15113 |
| Collagen formation | 4.30E-04 | 4.30E-04 | O60568;P02462;Q99715;Q15113 |
| Diseases associated with glycosaminoglycan metabolism | 1.03E-03 | 1.03E-03 | P13611;P07686;P51884 |
| Regulation of Insulin-like Growth Factor (IGF) transport and uptake by Insulin-like Growth Factor Binding Proteins (IGFBPs) | 1.04E-03 | 1.04E-03 | P09382;P35555;P13611;P08253 |
| Diseases of glycosylation | 1.70E-03 | 1.70E-03 | P13611;P35442;P07686;P51884 |
| Assembly of collagen fibrils and other multimeric structures | 2.01E-03 | 2.01E-03 | P02462;Q99715;Q15113 |
| HDL remodeling | 2.01E-03 | 2.01E-03 | P55058-1;P55058-2 |
| CS/DS degradation | 2.65E-03 | 2.65E-03 | P13611;P07686 |
| ECM proteoglycans | 2.65E-03 | 2.65E-03 | P02462;P13611;P51884 |
| Plasma lipoprotein assembly, remodeling, and clearance | 2.65E-03 | 2.65E-03 | P61916;P55058-1;P55058-2 |
| Keratan sulfate degradation | 2.70E-03 | 2.70E-03 | P07686;P51884 |
| Integrin cell surface interactions | 3.21E-03 | 3.21E-03 | P02462;P35555;P51884 |
| Crosslinking of collagen fibrils | 3.40E-03 | 3.40E-03 | P02462;Q15113 |
| Post-translational protein phosphorylation | 5.36E-03 | 5.36E-03 | P09382;P35555;P13611 |
| Diseases of metabolism | 5.80E-03 | 5.80E-03 | P13611;P35442;P07686;P51884 |
| Signaling by Nuclear Receptors | 7.14E-03 | 7.14E-03 | P55058-1;P07339;P08253;P55058-2 |
| Glycosaminoglycan metabolism | 7.14E-03 | 7.14E-03 | P13611;P07686;P51884 |
| Defective HEXB causes GM2G2 | 7.61E-03 | 7.61E-03 | P07686 |
| Plasma lipoprotein remodeling | 7.61E-03 | 7.61E-03 | P55058-1;P55058-2 |
| Metabolism of carbohydrates | 7.61E-03 | 7.61E-03 | P00505;P13611;P07686;P51884 |
| Keratan sulfate/keratin metabolism | 7.61E-03 | 7.61E-03 | P07686;P51884 |
| Molecules associated with elastic fibres | 8.03E-03 | 8.03E-03 | P23142;P35555 |
| NR1H3 & NR1H2 regulate gene expression linked to cholesterol transport and efflux | 8.47E-03 | 8.47E-03 | P55058-1;P55058-2 |
| Elastic fibre formation | 1.09E-02 | 1.09E-02 | P23142;P35555 |
| Collagen chain trimerization | 1.09E-02 | 1.09E-02 | P02462;Q99715 |
| NR1H2 and NR1H3-mediated signaling | 1.09E-02 | 1.09E-02 | P55058-1;P55058-2 |
| Chondroitin sulfate/dermatan sulfate metabolism | 1.09E-02 | 1.09E-02 | P13611;P07686 |
| **Signaling by PDGF** | 1.55E-02 | 1.55E-02 | P02462;P35442 |
| Defective B4GALT1 causes B4GALT1-CDG (CDG-2d) | 4.25E-02 | 4.25E-02 | P51884 |
| Defective CHSY1 causes TPBS | 4.25E-02 | 4.25E-02 | P13611 |
| Defective CHST14 causes EDS, musculocontractural type | 4.25E-02 | 4.25E-02 | P13611 |
| Defective CHST6 causes MCDC1 | 4.25E-02 | 4.25E-02 | P51884 |
| Defective CHST3 causes SEDCJD | 4.25E-02 | 4.25E-02 | P13611 |
| Defective ST3GAL3 causes MCT12 and EIEE15 | 4.25E-02 | 4.25E-02 | P51884 |
| Dermatan sulfate biosynthesis | 4.25E-02 | 4.25E-02 | P13611 |
| Hyaluronan uptake and degradation | 4.25E-02 | 4.25E-02 | P07686 |
| Aspartate and asparagine metabolism | 4.25E-02 | 4.25E-02 | P00505 |

| *Supplementary Table 14 – Name of the proteins associated with biological processes altered under low stiffness conditions* | | |
| --- | --- | --- |
| **Biological process** | **Accession number** | **Name** |
| ***Protein folding*** | P45877 | Peptidyl-prolyl cis-trans isomerase C |
|  | P10809 | 60 kDa heat shock protein, mitochondrial |
|  | P07237 | Protein disulfide-isomerase |
|  | P13667 | Protein disulfide-isomerase A4 |
|  | P50502 | Hsc70-interacting protein |
|  | P27797 | Calreticulin |
|  | P27824 | Calnexin |
|  | P23284 | Peptidyl-prolyl cis-trans isomerase B |
|  | Q15084 | Protein disulfide-isomerase A6 |
|  | P30101 | Protein disulfide-isomerase A3 |
| **Carbohydrate metabolic process** | P15121 | Aldo-keto reductase family 1 member B1 |
|  | P40926 | Malate dehydrogenase, mitochondrial |
|  | P30837 | Aldehyde dehydrogenase X, mitochondrial |
|  | P46926 | Glucosamine-6-phosphate isomerase 1 |
|  | P36222 | Chitinase-3-like protein 1 |
|  | O75390 | Citrate synthase, mitochondrial |
|  | Q04446 | 1,4-alpha-glucan-branching enzyme |
|  | Q14697 | Neutral alpha-glucosidase AB |
|  | P40925 | Malate dehydrogenase, cytoplasmic |
| **Tricarboxylic acid cycle** | P40926 | Malate dehydrogenase, mitochondrial |
|  | P07954 | Fumarate hydratase, mitochondrial |
|  | O75390 | Citrate synthase, mitochondrial |
|  | P40925 | Malate dehydrogenase, cytoplasmic |
| **Glycolytic process** | P19367 | Hexokinase-1 |
|  | P60174 | Triosephosphate isomerase |
|  | P04075 | Fructose-bisphosphate aldolase A |
|  | P04406 | Glyceraldehyde-3-phosphate dehydrogenase |
| **Cell redox homeostasis** | P30048 | Thioredoxin-dependent peroxide reductase, mitochondrial |
|  | Q16881 | Thioredoxin reductase 1, cytoplasmic |
|  | Q13162 | Peroxiredoxin-4 |
|  | P00390 | Glutathione reductase, mitochondrial |
| **Translation** | P60866 | 40S ribosomal protein S20 |
|  | P30050 | 60S ribosomal protein L12 |
|  | P15880 | 40S ribosomal protein S2 |
|  | Q04637 | Eukaryotic translation initiation factor 4 gamma 1 |
|  | P62081 | 40S ribosomal protein S7 |
|  | P62249 | 40S ribosomal protein S16 |
|  | P08865 | 40S ribosomal protein SA |
|  | P62277 | 40S ribosomal protein S13 |
|  | P61353 | 60S ribosomal protein L27 |
|  | P46781 | 40S ribosomal protein S9 |
|  | P42677 | 40S ribosomal protein S27 |
|  | P62701 | 40S ribosomal protein S4, X isoform |
|  | P46783 | 40S ribosomal protein S10 |
|  | P25398 | 40S ribosomal protein S12 |

| *Supplementary Table 15 – Protein names associated to biological processes altered under physioxia environments (long exposure)* | | |
| --- | --- | --- |
| **Biological process** | **Accession number** | **Name** |
| ***Immune response*** | P01877 | Immunoglobulin heavy constant alpha 2 |
|  | P02747 | Complement C1q subcomponent subunit C |
|  | A0A0C4DH72 | Immunoglobulin kappa variable 1-6 |
|  | A0A075B6H7 | Probable non-functional immunoglobulin kappa variable 3-7 |
|  | P04004 | Vitronectin |
|  | A0A0C4DH25 | Immunoglobulin kappa variable 3D-20 |
|  | P01701 | Immunoglobulin lambda variable 1-51 |
|  | P01834 | Immunoglobulin kappa constant |
|  | P80748 | Immunoglobulin lambda variable 3-21 |
|  | P01880 | Immunoglobulin heavy constant delta |
|  | P06331 | Immunoglobulin heavy variable 4-34 |
|  | P01876 | Immunoglobulin heavy constant alpha 1 |
|  | P01743 | Immunoglobulin heavy variable 1-46 |
|  | P0DP09 | Immunoglobulin kappa variable 1-13 |
|  | P01602 | Immunoglobulin kappa variable 1-5 |
|  | P06312 | Immunoglobulin kappa variable 4-1 |
|  | P01611 | Immunoglobulin kappa variable 1D-12 |
|  | P07357 | Complement component C8 alpha chain |
|  | A0A075B6R9 | Probable non-functional immunoglobulin kappa variable 2D-24 |
|  | P01703 | Immunoglobulin lambda variable 1-40 |
|  | A0A087WSY6 | Immunoglobulin kappa variable 3D-15 |
|  | P04433 | Immunoglobulin kappa variable 3-11 |
|  | P01024 | Complement C3 |
| **Fibrinolysis** | P00734 | Prothrombin |
|  | P04196 | Histidine-rich glycoprotein |
|  | P07355 | Annexin A2 |
|  | P05155 | Plasma protease C1 inhibitor |
|  | P00748 | Coagulation factor XII |
|  | P00747 | Plasminogen |
|  | P07225 | Vitamin K-dependent protein S |
|  | P03952 | Plasma kallikrein |
| **Cell adhesion** | P05067 | Amyloid-beta precursor protein |
|  | Q14126 | Desmoglein-2 |
|  | Q15262 | Receptor-type tyrosine-protein phosphatase kappa |
|  | O43854 | EGF-like repeat and discoidin I-like domain-containing protein 3 |
|  | Q9BUD6 | Spondin-2 |
|  | O94985 | Calsyntenin-1 |
|  | P78539 | Sushi repeat-containing protein SRPX |
|  | Q08629 | Testican-1 |
|  | Q08380 | Galectin-3-binding protein |
|  | P55287 | Cadherin-11 |
|  | P15291 | Beta-1,4-galactosyltransferase 1 |
|  | P19022 | Cadherin-2 |
|  | O60462 | Neuropilin-2 |
|  | Q92859 | Neogenin |
|  | P35442 | Thrombospondin-2 |
|  | Q9Y6C2 | EMILIN-1 |
|  | P12109 | Collagen alpha-1(VI) chain |
|  | P12111 | Collagen alpha-3(VI) chain |
|  | Q14112 | Nidogen-2 |
|  | Q15582 | Transforming growth factor-beta-induced protein ig-h3 |
|  | P11047 | Laminin subunit gamma-1 |
|  | Q99715 | Collagen alpha-1(XII) chain |
|  | Q13308 | Inactive tyrosine-protein kinase 7 |
|  | P20908 | Collagen alpha-1(V) chain |
|  | P39060 | Collagen alpha-1(XVIII) chain |
|  | P13611 | Versican core protein |
|  | P02751 | Fibronectin |
| **Angiogenesis** | Q6UVK1 | Chondroitin sulfate proteoglycan 4 |
|  | O60687 | Sushi repeat-containing protein SRPX2 |
|  | Q16610 | Extracellular matrix protein 1 |
|  | P13521 | Secretogranin-2 |
|  | O60462 | Neuropilin-2 |
|  | P08572 | Collagen alpha-2(IV) chain |
|  | P08253 | 72 kDa type IV collagenase |
|  | Q15582 | Transforming growth factor-beta-induced protein ig-h3 |
|  | P39060 | Collagen alpha-1(XVIII) chain |
|  | P98160 | Basement membrane-specific heparan sulfate proteoglycan core protein |
|  | P02751 | Fibronectin |
| **Ossification** | O95633 | Follistatin-related protein 3 |
|  | P02452 | Collagen alpha-1(I) chain |
|  | Q16610 | Extracellular matrix protein 1 |
|  | Q9Y240 | C-type lectin domain family 11 member A |
|  | P55287 | Cadherin-11 |
|  | P05997 | Collagen alpha-2(V) chain |

| *Supplementary Table 16 – Name of the proteins mapped on the significant downregulated Reactome pathways of common proteins shared by mechanomodulation and physioxia (readapted)* | | |
| --- | --- | --- |
| **Reactome pathway** | **Accession number** | **Name** |
| ***Extracellular matrix organization*** | P01033 | Metalloproteinase inhibitor 1 |
|  | P02462 | Collagen alpha-1(IV) chain |
|  | P02461 | Collagen alpha-1(III) chain |
|  | P05997 | Collagen alpha-2(V) chain |
|  | Q92626 | Peroxidasin homolog |
|  | P35555 | Fibrillin-1 |
|  | P20908 | Collagen alpha-1(V) chain |
|  | P12111 | Collagen alpha-3(VI) chain |
|  | P11047 | Laminin subunit gamma-1 |
|  | P21810 | Biglycan |
|  | P08123 | Collagen alpha-2(I) chain |
|  | Q99715 | Collagen alpha-1(XII) chain |
|  | Q15113 | Procollagen C-endopeptidase enhancer 1 |
| **ECM proteoglycans** | P02462 | Collagen alpha-1(IV) chain |
|  | P02461 | Collagen alpha-1(III) chain |
|  | P05997 | Collagen alpha-2(V) chain |
|  | P21810 | Biglycan |
|  | P08123 | Collagen alpha-2(I) chain |
|  | P20908 | Collagen alpha-1(V) chain |
|  | P12111 | Collagen alpha-3(VI) chain |
|  | P11047 | Laminin subunit gamma-1 |
| **Signal transduction** | P07602 | Prosaposin |
|  | P02462 | Collagen alpha-1(IV) chain |
|  | P02461 | Collagen alpha-1(III) chain |
|  | P05997 | Collagen alpha-2(V) chain |
|  | P08123 | Collagen alpha-2(I) chain |
|  | P35555 | Fibrillin-1 |
|  | P20908 | Collagen alpha-1(V) chain |
|  | P08476 | Inhibin beta A chain |
|  | P35442 | Thrombospondin-2 |
|  | P02786 | Transferrin receptor protein 1 |
|  | P12111 | Collagen alpha-3(VI) chain |
|  | P11047 | Laminin subunit gamma-1 |
| **Signaling by PDGF** | P02462 | Collagen alpha-1(IV) chain |
|  | P02461 | Collagen alpha-1(III) chain |
|  | P05997 | Collagen alpha-2(V) chain |
|  | P20908 | Collagen alpha-1(V) chain |
|  | P35442 | Thrombospondin-2 |
|  | P12111 | Collagen alpha-3(VI) chain |
| **Signaling by MET** | P02461 | Collagen alpha-1(III) chain |
|  | P05997 | Collagen alpha-2(V) chain |
|  | P08123 | Collagen alpha-2(I) chain |
|  | P20908 | Collagen alpha-1(V) chain |
|  | P11047 | Laminin subunit gamma-1 |

| *Supplementary Table 17 – Name of the proteins mapped on the significant mechano-upregulated pathways and physioxia-downregulated Reactome pathways of common proteins* | | |
| --- | --- | --- |
| ***Calnexin/calreticulin cycle*** | Q14697 | Neutral alpha-glucosidase AB |
|  | P30101 | Protein disulfide-isomerase A3 |
|  | P27797 | Calreticulin |
| ***Glucose metabolism*** | P00505 | Aspartate aminotransferase, mitochondrial |
|  | P40926 | Malate dehydrogenase, mitochondrial |
